# Phylogenomics of the ecdysteroid kinase-like (EcKL) gene family in insects highlights roles in both steroid hormone metabolism and detoxification

**DOI:** 10.1101/2023.06.29.546846

**Authors:** Jack L. Scanlan, Charles Robin

## Abstract

The evolutionary dynamics of large gene families can offer important insights into the functions of their individual members. While the ecdysteroid kinase-like (EcKL) gene family has previously been linked to the metabolism of both steroid moulting hormones and xenobiotic toxins, the functions of nearly all EcKL genes are unknown and there is little information on their evolution across all insects. Here, we perform comprehensive phylogenetic analyses on a manually annotated set of EcKL genes from 140 insect genomes, revealing the gene family is comprised of at least 13 subfamilies that differ in retention and stability. Our results show the only two genes known to encode ecdysteroid kinases belong to different subfamilies and therefore ecdysteroid metabolism functions must be spread throughout the EcKL family. We also provide comparative phylogenomic evidence that EcKLs are involved in detoxification across insects, with positive associations between family size and dietary chemical complexity, and also find similar evidence for the cytochrome P450 and glutathione S-transferase gene families. Unexpectedly, we find that the size of the clade containing a known ecdysteroid kinase is positively associated with host plant taxonomic diversity in Lepidoptera, possibly suggesting multiple functional shifts between hormone and xenobiotic metabolism. This work provides a robust framework for future functional studies into the EcKL gene family and opens promising new avenues for exploring the genomic basis of dietary adaptation in insects, including the classically studied co-evolution of butterflies with their host plants.

**Significance statement:** The ecdysteroid kinase-like (EcKL) gene family has been linked to both steroid inactivation and detoxification in insects, but the functions of most of its members are unknown. Here, we study the evolutionary history of the EcKLs and showcase how phylogenetics can inform the functional characterisation of enzyme families. EcKL family size varies over an order of magnitude and is associated with chemically complex diets, implicating numerous genes in detoxification. We also find a surprising link between a steroid-metabolising clade and host plant diversity in butterflies and moths, suggesting new detoxification functions may evolve through an insect-plant ‘chemical arms-race’. This work advances numerous functional hypotheses for multiple EcKL clades and proposes a new classification system for this poorly characterised gene family.

## Introduction

Variation in large multigene families (hereafter ‘gene families’) is thought to underpin many aspects of phenotypic diversity across the tree of life (Lespinet et al. 2002; Demuth & Hahn 2009; Hahn et al. 2007). However, for most gene families, functional knowledge is limited to only a handful of their members, leaving most genes uncharacterised; this problem will only grow larger as more and more genomes are sequenced (Niehaus et al. 2015; Nagy et al. 2020; Ellens et al. 2017).

A powerful method for predicting gene function is phylogenomics, which uses the evolutionary history of a gene to understand its functional constraints and taxonomic distribution (Eisen 1998). This is especially useful in gene families, where processes like gene duplication, neofunctionalisation, subfunctionalisation and pseudogenisation produce complex gene-level phylogenies of orthologs and paralogs through a ‘birth-death’ process (Nei & Rooney 2005; Prince & Pickett 2002). Ascribing phenotypic traits to evolutionary properties of specific genes or clades within gene families can produce functional hypotheses to test through experimental methods. Functional phylogenomic approaches have been fruitfully applied to gene families across a wide range of taxa (Delaux et al. 2014; Finet et al. 2019; Li & Zhang 2014). The most statistically robust of these studies use phylogenetic comparative methods to account for trait correlations due to evolutionary relationships (Uyeda et al. 2018; Cornwell & Nakagawa 2017).

Three evolutionary properties of genes that are commonly of interest in phylogenomics are: (1) sequence conservation—nucleotide or amino acid identity between homologs; (2) presence conservation—presence or absence of specific genes in taxa of interest (also called gene retention); and (3) copy-number conservation—tendency of orthologs to remain single-copy rather than multi-copy (also called duplicability; Waterhouse et al. 2011; Cooper & Brown 2008). These types of conservation vary with the function of particular gene family clades—for example, enzymes that show high presence and copy-number conservation are more likely to be involved in essential biosynthetic processes than lineage-specific xenobiotic metabolism (Tan & Low 2018; Thomas 2007; Kawashima & Satta 2014).

Metabolic detoxification, a subset of xenobiotic metabolism, is a biochemical and physiological process that allows organisms to tolerate toxic compounds present in their immediate environment. Knowledge of detoxification processes in insects is essential for understanding their ecology, co-evolution with other taxa, insecticide resistance in pest species, and insecticide sensitivity in beneficial species (Alyokhin & Chen 2017; Arena & Sgolastra 2014; De Fine Licht et al. 2013; Simon et al. 2015). Multiple gene families, encoding both enzymes and transporters, contribute to detoxification in eukaryotes, including cytochrome P450s (P450s), glutathione S-transferases (GSTs), carboxyl/cholinesterases (CCEs), UDP-glycosyltransferases (UGTs) and ABC transporters (ABCs; Omiecinski et al. 2011). All of these gene families exist in insects and have been associated with tolerance of, or resistance to, toxins such as phytochemicals, microbial secondary metabolites or synthetic insecticides (Li et al. 2007). The sizes and composition of detoxification gene families are thought to evolve in response to changes in ecologically relevant exposure to xenobiotic compounds, although tests of this hypothesis on a broad scale across insects have been limited (Rane et al. 2019, 2016; Calla et al. 2017; Breeschoten et al. 2021).

Many aspects of the physiology and biochemistry of detoxification in insects have been well studied (Li et al. 2007; Yang et al. 2007; Dow et al. 2018; Wilkinson 1986), but others are poorly understood. A notable example of the latter is xenobiotic phosphorylation—the addition of negatively charged phosphate groups to hydroxyl moieties to xenobiotic compounds or their metabolites—a reaction that occurs widely in insects but rarely in other animals (Mitchell 2015). Phosphorylated metabolites of phenolic, steroidal and glycosidic compounds have been identified in a taxonomically broad range of insects (eg. Easson et al. 2021; Olsen et al. 2015, 2014; Boeckler et al. 2016; reviewed in Scanlan et al. 2020), suggesting phosphorylation is ubiquitous and potentially important across all insects. Despite this, kinases acting on specific xenobiotic compounds (‘detoxicative kinases’) have, to the best of our knowledge, yet to be identified in any insect species.

Ecdysteroids are polyhydroxylated steroid hormones that control insect development, physiology and behaviour, and are synthesised from dietary sterols (Feldlaufer et al. 1995; Carlisle & Jenkin 1959; Yamanaka et al. 2013). In most insects, the key ecdysteroid is typically considered 20-hydroxyecdysone (20E), although the precursor ecdysone (E) may also have specific functions (Ono 2014; Riddiford 1993). Active ecdysteroid titres can be regulated both through expression and activity of their biosynthetic enzymes (encoded by the ‘Halloween’ genes; Rewitz et al. 2006) and through catabolism/inactivation via various modification or conjugation reactions (Rees 1995; Rharrabe et al. 2007). An important ecdysteroid conjugation reaction is phosphorylation, which can occur at the C-2, C-3, C-22 or C-26 positions on the ecdysteroid nucleus. Ecdysteroid phosphorylation is reversible (Davies et al. 2007; Yamada & Sonobe 2003) and thus plays important roles in hormone storage and recycling during insect development and reproduction (Sonobe & Ito 2009; Rees & Isaac 1984). It is mediated by ecdysteroid kinases, of which only two have been both genetically and biochemically characterised (Sonobe et al. 2006; Peng et al. 2022). The diversity of ecdysteroid-phosphate conjugates found in insects (reviewed in Lafont et al. 2012) strongly suggests more ecdysteroid kinases are yet to be identified.

Both characterised ecdysteroid kinases belong to the ecdysteroid kinase-like (EcKL) gene family (InterPro entry IPR004119; Blum et al. 2021). EcKL genes are predicted to encode cytosolic kinases with small molecular substrates and are generally poorly functionally characterised, with a handful of exceptions. The two biochemically characterised EcKLs are *BmEc22K* in the silkworm *Bombyx mori* (Lepidoptera), which encodes an ecdysteroid 22-kinase involved in reproduction and early embryonic development (Fujinaga et al. 2020; Sonobe et al. 2006), and *AgEc2K* in the mosquito *Anopheles gambiae* (Diptera), which encodes a 20E 22-kinase that limits oviposition in blood-fed virgin females (Peng et al. 2022). *AgEcK1*, another EcKL in *A. gambiae,* may function in male survival but has not been studied biochemically (Peng et al. 2022). *JhI-26* is an EcKL in *Drosophila melanogaster* (Diptera) that responds to juvenile hormone (JH) signalling (Dubrovsky et al. 2000) and may play a role in *Wolbachia*-mediated cytoplasmic incompatibility (Liu et al. 2014). *Ipi10G08* (Genbank: AY875646.1) is a male-biased EcKL in the pine engraver beetle *Ips pini* (Coleoptera) that is also strongly inducible by JH (Bearfield et al. 2008). Lastly, *CHKov1* and *CHKov2* are EcKLs in *D. melanogaster* implicated in resistance to sigma virus infection through an unknown molecular mechanism (Magwire et al. 2011, 2012; Duxbury et al. 2019).

We previously hypothesised that other members of the EcKL gene family might encode the detoxicative kinases of insects (Scanlan et al. 2020). Our first test of this hypothesis, through the integration of transcriptomic, evolutionary and genetic association data in *Drosophila*, suggested that nearly half of EcKLs in *D. melanogaster* have characteristics common to known detoxification genes (Scanlan et al. 2020). Recently, functional genetic experiments further supported the detoxification hypothesis by implicating multiple EcKLs in the *Drosophila* clade Dro5 in tolerance to the developmental toxins caffeine and kojic acid (Scanlan et al. 2022). However, it is currently unclear if these genes function in detoxification in insects outside of *D. melanogaster*.

Here, we conduct comprehensive phylogenomic analyses of the EcKL gene family across insects and propose a working classification system for individual genes based on subfamilies and taxon-specific clades. We also conduct phylogenetic comparative analyses of EcKLs and known detoxification gene families (P450s, GSTs and CCEs) in relation to measures of exposure to xenobiotic diversity and further test the hypothesis that EcKLs encode detoxicative kinases. We aim to generate testable hypotheses for the functions to specific EcKL genes and clades, and to establish an evolutionary framework to aid the characterisation of novel ecdysteroid kinase enzymes in insects.

## Results

### The EcKL gene family is taxonomically limited to Tetraconata (Hexapoda + Crustacea) within Arthropoda

In gene family databases such as InterPro (Blum et al. 2021), the vast majority of EcKL sequences belong to arthropods. To rigorously explore the distribution of this gene family within the arthropods, we used BLAST searches to find EcKL sequences in NCBI transcriptome shotgun assemblies (TSAs) from the major lineages of arthropods spanning the four subphyla: Hexapoda (insects and allies), Crustacea (crustaceans), Myriapoda (centipedes and millipedes) and Chelicerata (spiders and allies). Numerous EcKL transcripts were found in hexapods (Insecta, Diplura, Collembola and Protura) and crustaceans, but in Chelicerata and Myriapoda, the evidence for the presence of the EcKL gene family was very poor (Fig. 1A). We found only 25 hits in TSAs from 1,216 species in Chelicerata (all in Arachnida). When reciprocally used as queries against the Arthropoda protein database, these sequences had very high identity to insect EcKLs—strongly suggesting they were due to RNA contamination from dietary insects—or had very low sequence similarity to EcKLs and when examined with InterProScan (Jones et al. 2014) belonged to a different gene family (‘Uncharacterised oxidoreductase Dhs-27’, IPR012877). As such, these sequences were not considered genomically encoded EcKLs belonging to chelicerates. We found a single EcKL-containing transcript (Genbank: GERS01005135.1) in Myriapoda, from a millipede in the genus *Craspedosoma*, whose highest sequence identity (40-47%) was to EcKLs in collembolan TSAs. However, reciprocal BLAST searches against the NCBI Myriapoda protein nr and nucleotide nt databases (spanning nine genome assemblies) using GERS01005135.1 as a query produced no significant hits, suggesting this single transcript is likely due to RNA contamination from an ingested, unknown hexapod. Overall, these data strongly suggest the EcKL gene family is limited, within Arthropoda, to Hexapoda + Crustacea, a clade commonly referred to as Tetraconata (Richter 2002; Schwentner et al. 2018).

**Figure 1.**
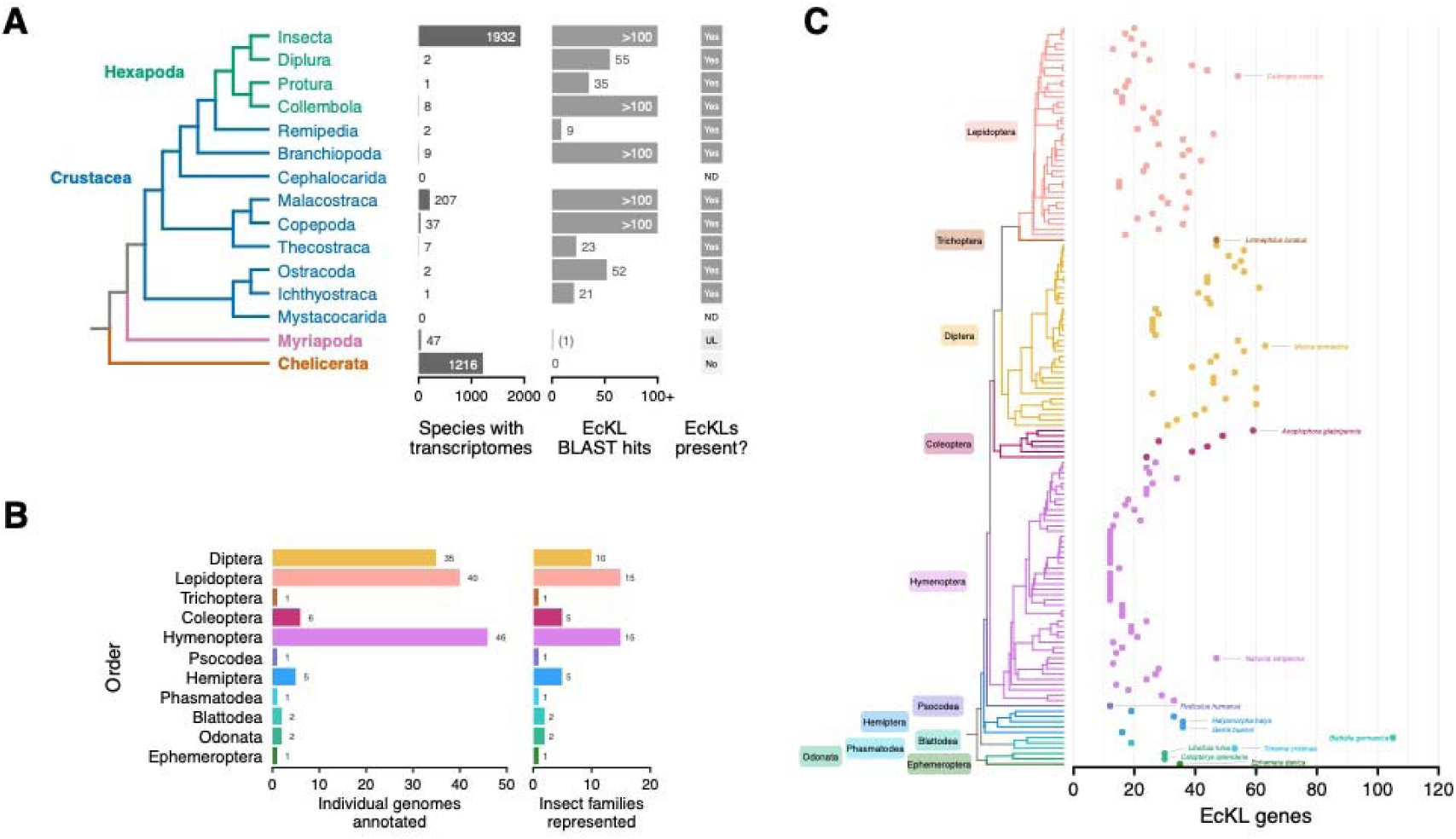
Identification and annotation of the EcKL gene family in 140 insect genomes. **A.** EcKLs are only reliably found in the transcriptomes of arthropods from the subphyla Hexapoda (green) and Crustacea (blue), based on tblastn searches in NCBI. The single EcKL found in a myriapod transcriptome (in brackets) is unlikely to be present in the genome; see text for details. NCBI BLAST search results are capped at 100 hits. Cladogram is derived from Schwentner et al. (2017) and is coloured by subphyla (subphyla names are bold). ND, not determined; UL, unlikely. **B.** EcKLs annotated in 140 insect genome assemblies spread across 11 taxonomic orders (left) and 59 taxonomic families (right). See Table S1 for details of the assemblies used. **C.** The number of EcKL genes annotated in insect genome assemblies varies between and within orders. For each order, the species with the largest number of EcKLs is labelled. The phylogenetic tree with species names and node age evidence is available graphically in Fig. S6, as well as in Newick file format in the Supplementary Data. Gene counts per species are available in Table S1.

For the remainder of this study, we focused on annotating and phylogenetically characterising EcKLs in insects alone, due to the large number of genomes available and the pre-existing functional focus on insect EcKLs in the published literature.

### Annotation of the EcKL gene family in 140 insect genome assemblies

The EcKL gene family was manually annotated in 140 insect genome assemblies, representing 139 species from 107 genera, 58 families, 43 superfamilies and 11 orders. Three orders were best represented: Hymenoptera (n = 46), Lepidoptera (n = 40) and Diptera (n = 35; Fig. 2B). In total, 4,214 EcKL gene models were annotated, comprising 3,905 (92.7%) full gene models (with one or more complete EcKL domain and plausible start and stop codons) and 305 (7.2%) partial gene models (containing an incomplete EcKL domain and/or an unknown start or stop codon; Table S1). The mean size of translations from full gene models was 418 aa, while the median was 414 aa. The frequency of deviations from single-domain protein architecture was low: 33 (0.8%) gene models encoded proteins with two predicted EcKL domains, while 17 (0.4%) gene models encoded proteins with additional N- or C-terminal disordered regions.

**Figure 2.**
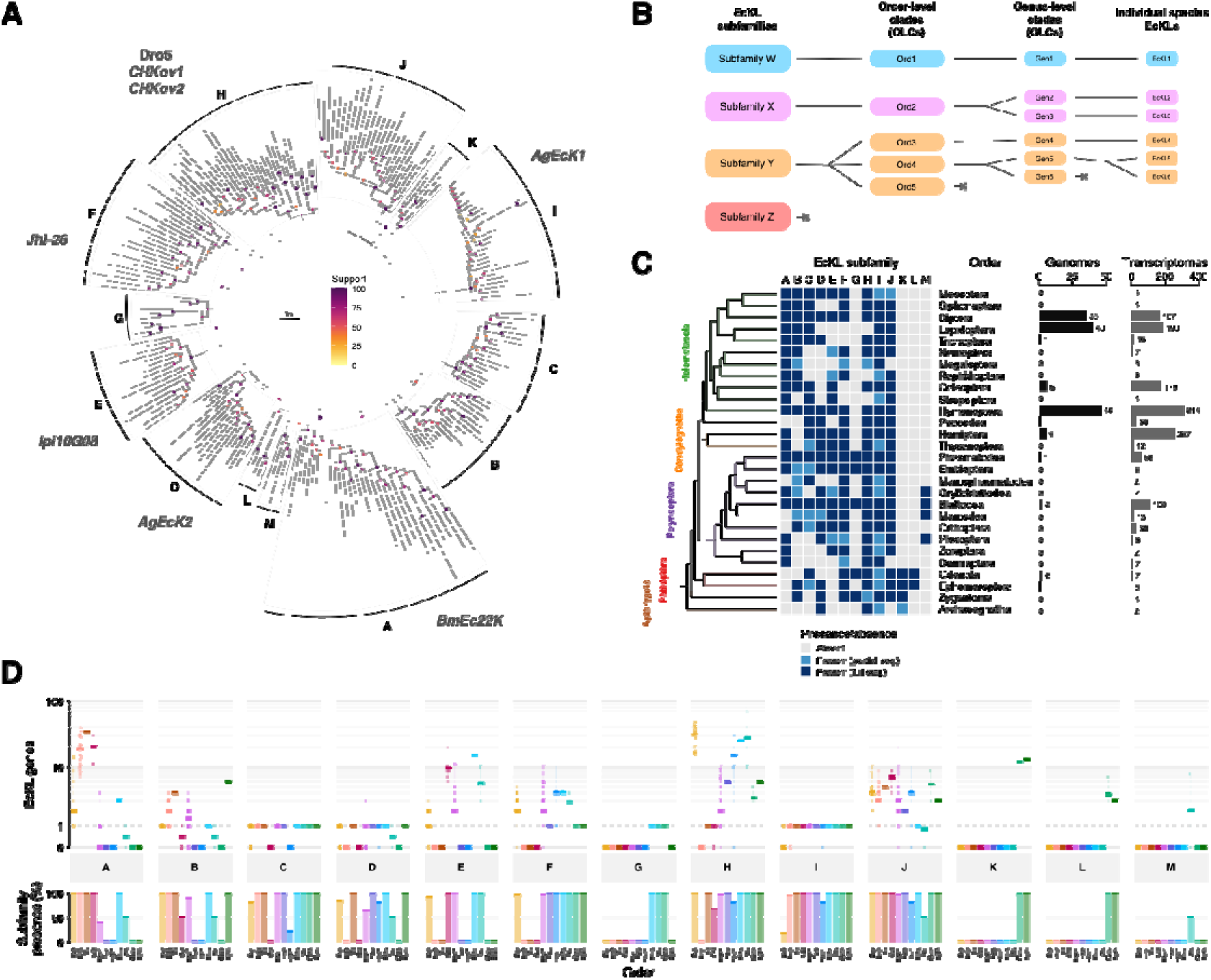
EcKL genes in insects can be classified into one of 13 ‘subfamilies’, from A to M. **A.** Unrooted ML phylogenetic tree derived from a dataset of [x] EcKL amino acid sequences from all 28 insect orders using IQ-TREE (Nguyen et al. 2015), defining the 13 subfamilies (shaded grey clades). Functionally characterised EcKLs belonging to specific subfamilies have been indicated in grey text; see text for details. Support values are derived from UFBoot2, where values ≥95 are considered reliable (Hoang et al. 2018); nodes with support ≥95 are slightly larger. The phylogenetic tree with tip labels is available in the Newick file format in the Supplementary Data. **B.** Overview of the working classification system for the EcKL gene family. Individual EcKLs can simultaneously belong to three groupings— subfamilies, order-level clades (OLCs) and genus-level clades (GLCs)—which are defined as clades present as a single gene in the most recent common ancestor of Insecta, the species’ order and the species’ genus, respectively. Illustrative examples of relationships between the groupings for hypothetical subfamilies (‘W’ through ‘Z’) are shown. Higher-level groups can contain single lower-level groups for single-copy orthologs (blue) or multiple lower-level groups due to gene duplication (pink and orange). Individual species may be missing EcKLs from any of the groupings due to gene deletion (orange and red). GLCs for the genus *Drosophila* were defined in Scanlan et al. (2020). **C.** Presence and absence of EcKL subfamilies in the 28 insect orders based on genomic and transcriptomic evidence. Genome numbers (black bars) indicate the number of genome assemblies annotated in this study (see Fig. 1B); transcriptome numbers (grey bars) indicate the number of TSAs in NCBI searched using tblastn. Phylogenetic tree is derived from Misof et al. (2014). **D.** EcKL subfamily gene count and presence conservation per annotated genome and order. Top: EcKL gene number in each subfamily for each of the 140 annotated genomes (dots) in 11 insect orders (colours). Solid lines indicate the median gene count per order and the range is depicted using a faint vertical line; note the log scale. A gene count of 1 for a particular subfamily is emphasised with a dotted line. Bottom: EcKL subfamily presence per insect order expressed as a percentage of genomes that contain at least 1 EcKL in that subfamily.

The number of EcKL genes per genome (mean = 30, median = 26) ranged substantially between species, from 12 in the human louse *Pediculus humanus* (Psocodea), Jerdon’s jumping ant *Harpegnathos saltator* (Hymenoptera) and 12 bee species (Hymenoptera), to 105 in the German cockroach *Blattella germanica* (Blattodea; Fig. 2C).

### Insect EcKLs can be classified into 13 subfamilies based on a phylogenetic analysis across all 28 insect orders

The EcKLs currently lack a standardised gene nomenclature like those of other enzyme families, such as the cytochrome P450s, UDP-glycosyltransferases and the glutathione S-transferases (references in Ahn et al. 2012; Enayati et al. 2005; Nelson 2006). We wished to develop a working, cladistic classification system for the EcKLs (Fig. 2B), not to act as a nomenclature *per se*, but to inform the functional characterisation of the gene family. Here we propose a scheme where subfamilies are highest level of classification and define them as clades derived from single genes present in the most recent common ancestor (MCRA) of all insects. We then use order-level clades (OLCs) and genus-level clades (GLCs) for lower levels of classification and while these are arbitrary, they are likely useful for most focused purposes. OLCs are clades derived from single genes present in the MCRA of a specific order, while GLCs are clades derived from single genes present in the MCRA of a specific genus. GLCs for the genus *Drosophila* (Ephydroidea: Diptera) were defined previously (Scanlan et al. 2020). These levels of classification exist in a hierarchy such that individual EcKL genes belong to a GLC, an OLC and a subfamily simultaneously (Fig. 2B).

Preliminary phylogenetic analyses of our entire dataset of annotated EcKLs suggested the existence of deep clades consistent with subfamilies. To help define these deeply branching parts of the EcKL phylogeny with a more tractable and evenly sampled dataset, we curated a collection of 241 EcKL sequences (‘core set’) belonging to putative subfamilies from all 28 insect orders (Misof et al. 2014); in insect orders where we had not annotated a genome, we searched NCBI TSAs using tblastn with putative subfamily members as queries. Maximum likelihood treebuilding with IQ-TREE (Nguyen et al. 2015) produced a gene phylogeny from which 13 subfamilies (A—M) were identified (Fig. 2A). Branch support for these subfamilies was generally high, but relatively low support at the base of some subfamilies (D, G, H and L) suggest these clades should be considered tentative and should be revisited in future studies with greater sampling of early diverging insect orders.

To explore the presence-conservation of EcKL subfamilies across insect orders, we classified all genomically annotated EcKLs into each subfamily by aligning each order’s sequences to our core set of EcKLs and generating phylogenies. We also used the core set as tblastn queries against TSAs from each insect order and, for each EcKL subfamily, classified the highest-identity hits as members or non-members of the subfamily by examination of phylogenies built from alignments of the hits and the core set. Together, this produced a presence-absence matrix for each subfamily by each order (Fig. 2C). One caveat for this approach is that querying transcriptomes likely has a high false negative rate. Despite this, this approach identified subfamilies I and J as present in all and nearly all orders, respectively, with J only missing from Megaloptera (n = 3 TSAs) and Dermaptera (n = 7 TSAs). The most taxonomically restricted subfamilies were G, K, L and M, which were missing from most orders, especially in Holometabola (Fig. 2C).

Next we focused on the 11 insect orders for which we had genomically annotated EcKLs to explore differences in presence-conservation and size of EcKL subfamilies (Fig. 2D). Subfamilies I and J were present in at least one species in all 11 orders, but were not always present within each species. Subfamily gene number varied substantially—the highest gene count for subfamily within a single genome was 75 in subfamily H for *Blattella germanica* (Blattodea), while the highest median subfamily size for a particular order (n ≥ 2 genomes) was 31 for subfamily H in Diptera. Subfamilies C, D, G and I tended to be conserved as single-copy orthologs, if present, while most other subfamilies showed substantial variation in copy number within and between orders (Fig. 2D). These data suggest the EcKL gene family has undergone dynamic changes in size between taxa, due to both gene duplication and loss.

Notably, pairwise amino acid sequence identities for the full-length EcKLs in our core set suggested the identity thresholds (for example the 40% for families and 55% as used for the P450s ; Nelson 2006), poorly capture the cladistic structure of the EcKLs. Eleven of the thirteen EcKL subfamilies had a minimum within-subfamily identity of <30% (Fig. S1). Most subfamilies had a minimum between-subfamily identity of <20%, indicating high levels of sequence divergence within the EcKL family in insects.

### Mapping known functions to EcKL subfamilies

Identification of EcKL subfamilies allowed us to place genes of known function within a phylogenetic context (Fig. 2A). The ecdysteroid 22-kinases *BmEc22K* and *AgEcK2* belong to subfamilies A and D, respectively, suggesting either ecdysteroid kinase activity evolved separately in these subfamilies, or the ancestral substrates for clades A, B, C, D, L and M are ecdysteroids (Fig. 2A). Curiously, while biochemically characterised ecdysteroid kinases are known in Orthoptera (Kabbouh & Rees 1991, 1993), this order may lack subfamily A, based on 30 TSAs (Fig. 2C); this makes subfamily D genes the most likely candidates to encode these enzymes. Two JH-responsive EcKLs, *JhI-26* in *D. melanogaster* and *Ipi10G08* in *I. pini*, belong to subfamilies F and E, respectively, which form a deep clade with subfamily G (Fig. 2A); these results could suggest a hypothesis that EcKLs from these subfamilies might be involved with JH signalling across various insects. *CHKov1* and *CHKov2*, genes associated with sigma virus resistance in *D. melanogaster*, as well as most of the genes suggested to be involved in detoxification in *D. melanogaster* in Scanlan et al. (2020) and Scanlan et al. (2022), belong to subfamily H. Lastly, *AgEcK1*, which may play a role in male survival in *A. gambiae*, belongs to the highly conserved subfamily I.

### Defining OLCs for Diptera, Lepidoptera and Hymenoptera

The availability of genomes in the orders Diptera (true flies, mosquitoes and allies), Lepidoptera (butterflies and moths) and Hymenoptera (bees, wasps, ants and sawflies) gave us sufficient resolution to infer OLCs for these taxa.

In Diptera, we inferred the existence of 20 OLC (Dip1–Dip20; Fig. S2), with only one clade, Dip12 (subfamily B) retained as single-copy orthologs in all 35 genomes. Dip1, Dip3, Dip4 and Dip7 (subfamily H) and Dip8 (subfamily A) have undergone substantial increases in size in some taxa, the most extreme example being 28 Dip4 genes in the housefly, *Musca domestica* (Fig. 3A).

**Figure 3.**
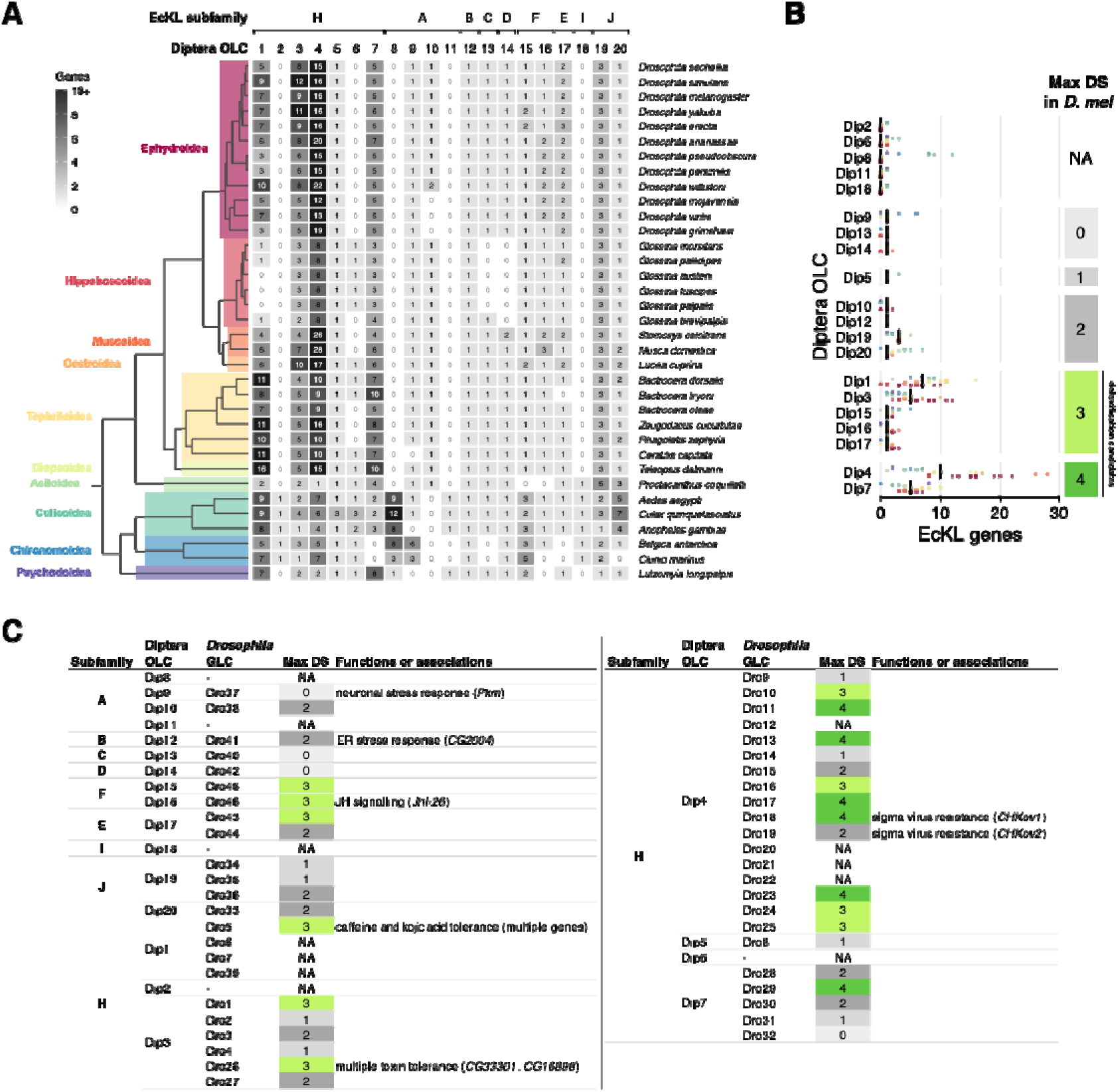
EcKL order-level clades (OLCs) for the order Diptera (true flies). **A.** Diptera OLCs for 35 species across 10 taxonomic superfamilies. Boxes show the number of genes per OLC per species (indicated by the shading intensity and the number inside). EcKL subfamily groupings per OLC are shown at top. See Fig. S2 for a tree of Diptera OLCs. The phylogenetic tree is derived from Fig. S6; coloured boxes indicate the superfamilies. **B.** Diptera OLCs gene counts per genome, grouped by the maximum ‘detoxification score’ (DS) for [component] *D. melanogaster* EcKLs as per Scanlan et al. (2020), where genes with DS ≥3 are considered detoxification candidates; ‘NA’ indicates an OLC missing from *D. melanogaster*. Vertical black lines are the median gene counts per OLC; dots are coloured by superfamily as per Fig. 3A. **C.** Mapping of Diptera OLCs and *Drosophila* GLCs to each other and the EcKL subfamilies, with additional max DS values (see Fig. 3B) and functional data in *D. melanogaster* (see text for references).

In Lepidoptera, we inferred the existence of 17 OLC (Lep1–Lep17; Fig. S3), with only one clade, Lep9 (subfamily C), retained as single-copy orthologs in all 40 genomes. Lep1, Lep3 and Lep8 (subfamily A) showed the largest expansions within this order; notably, the ecdysteroid 22-kinase *BmEc22K* belongs to the Lep1 clade and is the only member of this clade in *B. mori*.

In Hymenoptera, we inferred the existence of 22 OLC (Hym1–Hym22; Fig. S4), with two clades, Hym19 (subfamily I) and Hym21 (subfamily C), conserved as single-copy orthologs in all 46 genomes. Hym1 (subfamily H), Hym14 (subfamily F) and Hym17 (subfamily E) have expanded the most in this order, with 18 Hym1 genes in the fungus-growing ant *Trachymyrmex cornetzi* (Fig. S5).

### Mapping EcKL functions to OLCs across Diptera

As we previously assigned EcKLs in the *Drosophila* genus (superfamily Ephydroidea) to 46 GLCs (Dro1—Dro46; Scanlan et al. 2020), we mapped functions from EcKLs in the model insect *D. melanogaster* back to OLC in Diptera. One key functional metric is the ‘detoxification score’, which indicates how likely a gene is to be involved in metabolic detoxification in *D. melanogaster* by integrating phylogenetic stability, transcriptional induction and tissue-specific expression, where a DS of 3 or higher is the threshold for detoxification candidacy (Scanlan et al. 2020). Notably, some Diptera OLCs that contain *D. melanogaster* detoxification candidate genes have some of the highest median gene counts across the order, especially Dip1, Dip3, Dip4 and Dip7 (all subfamily H), which have substantially expanded in most dipterans (Fig. 3B). Dip1 contains the Dro5 EcKLs, which have been experimentally linked to caffeine and kojic acid tolerance (Scanlan et al. 2022), while Dip3 contains two close paralogs that have been associated with tolerance to ethanol, imidacloprid and/or methylmercury (Scanlan et al. 2020; Fig. 3C). These data suggest EcKL clade expansions in Diptera might be tied in many cases to detoxification, and Dip1, Dip3 and Dip4 EcKLs in other dipterans should be investigated for possible detoxification functions.

*AgEcK2*, which encodes a 20E 22-kinase in the mosquito *A. gambiae* (Peng et al. 2022), belongs to subfamily D and therefore the Diptera OLC Dip14. This clade is broadly conserved as single-copy orthologs across Diptera, except for in the *Glossina* genus of tsetse flies, where it has been lost, and the stable fly *Stomoxys calcitrans*, where it is present as two paralogs (Fig. 3A). This high level of conservation suggests the *D. melanogaster* Dip14 gene *CG5644* may also encode an ecdysteroid kinase.

### EcKL family size is positively associated with increasing dietary chemical complexity across insects

We wished to use our dataset of genomically annotated genes in 140 species to test for detoxification functions of EcKLs across all insects, with the hypothesis that species with higher levels of exposure to xenobiotic compounds would have a larger number of genes. Additionally, we used overlapping data from Rane et al. (2019) for the sizes of the P450, GST and CCE gene families (in 98 species) to explore their relationships to our measures of detoxification ability. We classified the 140 annotated insect species into one of five dietary categories (detritivorous, carnivorous, herbivorous, haematophagous and pollen-feeding), and into one of three ‘xenobiotic diversity’ (XD) levels (small, medium, large) based on estimates of their ecologically relevant exposure to xenobiotic compounds. We then fit phylogenetically aware linear models using a dated time tree (Fig. S6) and phylogenetic least-squares regression (PGLS) to explain gene family size using various predictor variables.

We first found that the gene family sizes of EcKLs, P450s, GSTs and CCEs are all significantly positively associated with each other, with R^2^ between 0.053–0.074 (EcKLs and CCEs) and 0.283–0.306 (P450s and GSTs; Fig. S7, Table S2). While this is consistent with our hypothesis, it could be confounded by other factors such as total gene content, genome size or relaxed selection on gene duplication, and caution should be taken interpreting these results.

Diet was a significant predictor of gene family size for the EcKLs (F_4,135_ = 7.56, p < 1.0×10^-4^), P450s (F_4,93_ = 4.63, p = 0.0019) and GSTs (F_4,93_ = 6.56, p < 1.0×10^-4^), but not the CCEs (F_4,93_ = 1.28, p = 0.28; Fig. 4A–D, Table S3). Pairwise comparisons revealed the largest and most consistent mean differences were between haematophagous insects and detritivorous insects; there were no significant differences between pollen-feeding insects and other groups for any gene family, despite some large effect sizes in the EcKLs and P450s (Fig. S8).

**Figure 4.**
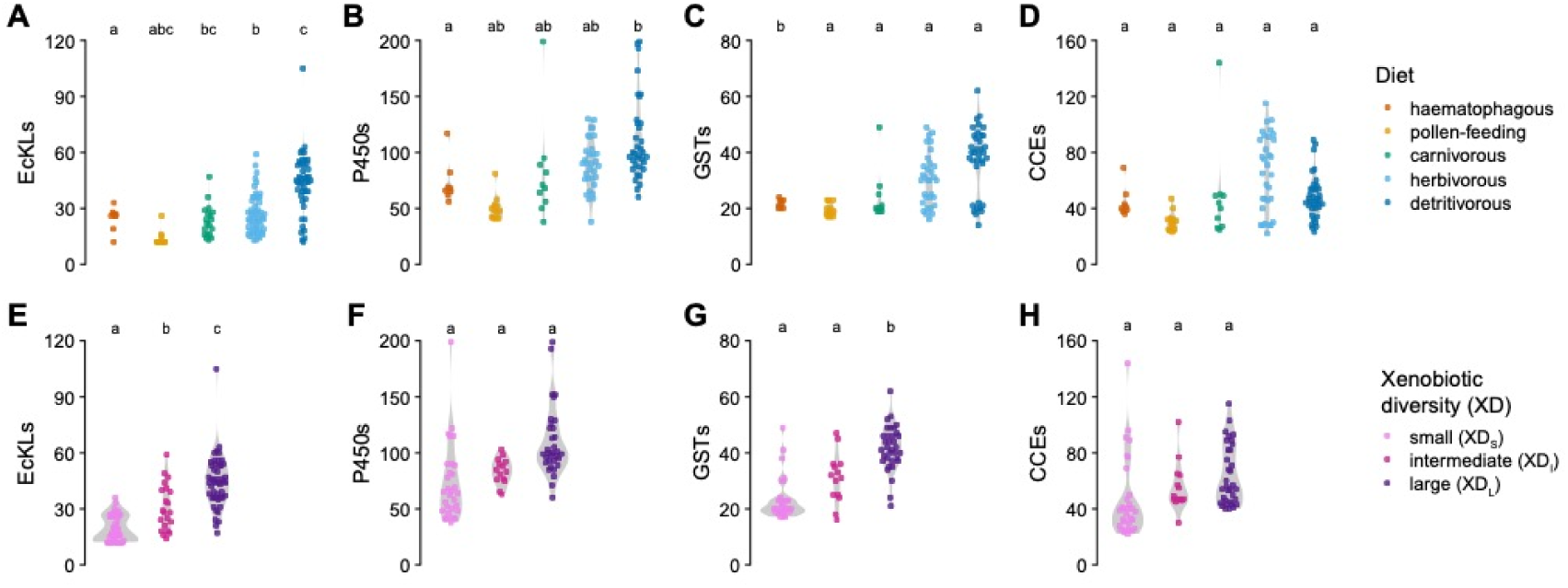
Differences in gene family size between dietary categories (A–D) and an estimate of xenobiotic diversity (XD; E–H) for the EcKLs (A,E), P450s (B,F), GSTs (C,G) and CCEs (D,H). Data are displayed as dots (insect species) and violins (distribution within each group). Groups that do not share a letter within each plot have significantly different means (adjusted p < 0.05) based on PGLS regression. **A–D.** For effect sizes, see Fig. S8; for model summaries, see Table S3. **E–H.** Data shown is a subset (n = 115 for EcKLs, n = 79 for P450s, GSTs and CCEs) with taxa with ambiguous XD estimates removed; for effect sizes of analyses run on the full dataset (n = 140 for EcKLs, n = 98 for P450s, GSTs and CCEs), including effect sizes, see Fig. S9; for model summaries, see Table S4.

As XD assignment for some taxa (ants and wasps) was ambiguous, we conducted analyses of XD as predictor of gene family size using two datasets, once with the full dataset (n = 140 species for EcKLs, n = 98 species for P450s/GSTs/CCEs) and once with a subset with those taxa removed (n = 115 species for EcKLs, n = 78 species for P450s/GSTs/CCEs). In the full dataset, XD was a significant predictor of gene family size for the EcKLs (F_2,137_ = 29.4, p < 1.0×10^-4^), P450s (F_2,95_ = 3.51, p = 0.034) and GSTs (F_2,95_ = 9.99, p = 1.0×10^-4^), but not the CCEs (F_2,95_ = 0.161, p = 0.85; Table S4). In the dataset with some ants and wasps removed, the P450 prediction became non-significant (F_2,75_ = 1.44, p = 0.24), while the EcKL (F_2,112_ = 27.1, p < 1.0×10^-4^) and GST (F_2,75_ = 10.8, p = 1.0×10^-4^) predictions remained significant (Fig. 4E–H, Table S4). Pairwise comparisons showed significant effects were always in the hypothesised direction, with larger XD estimates always resulting in similar or higher gene family sizes (Fig. S9).

Overall, these data support our hypothesis that EcKL family size is associated with increasing dietary chemical complexity across insects, and also demonstrate similar results for the GSTs and P450s, but not the CCEs.

### EcKL family size is positively associated with host plant diversity in Lepidoptera

As a more focused test of EcKL function in detoxification, we used host plant data from the HOSTS database (Robinson et al. 2010) as a proxy for dietary xenobiotic diversity in herbivorous Lepidoptera, which are known to closely co-evolve with their host plants (Ehrlich & Raven 1964; Menken et al. 2010). Across the species with host plant data (n = 38), there was substantial variation in taxonomic diversity, from strict monophagy (one host species—*Bombyx mandarina*, *Bicyclus anynana* and *Calephelis virginiensis*) to extreme polyphagy (215 species from 166 genera and 72 families—*Spodoptera litura*; Fig. 5A). Our hypothesis was that the number of host plant species, genera and families associated with each lepidopteran species would positively predict the size of the EcKL gene family. We fit phylogenetically aware linear models using the Lepidoptera subset of our dated time tree (Fig. S6) and PGLS to explain EcKL family size, as well as the size of each Lepidoptera OLC (Fig. 5A), using log_2_-transformed counts of host species, genera and families as predictor variables (Table S5). We did not conduct similar analyses with the GST, P450 or CCE gene families due to the limited overlap (n = 16 species) between our annotations and those of Rane et al. (2019) in Lepidoptera.

**Figure 5.**
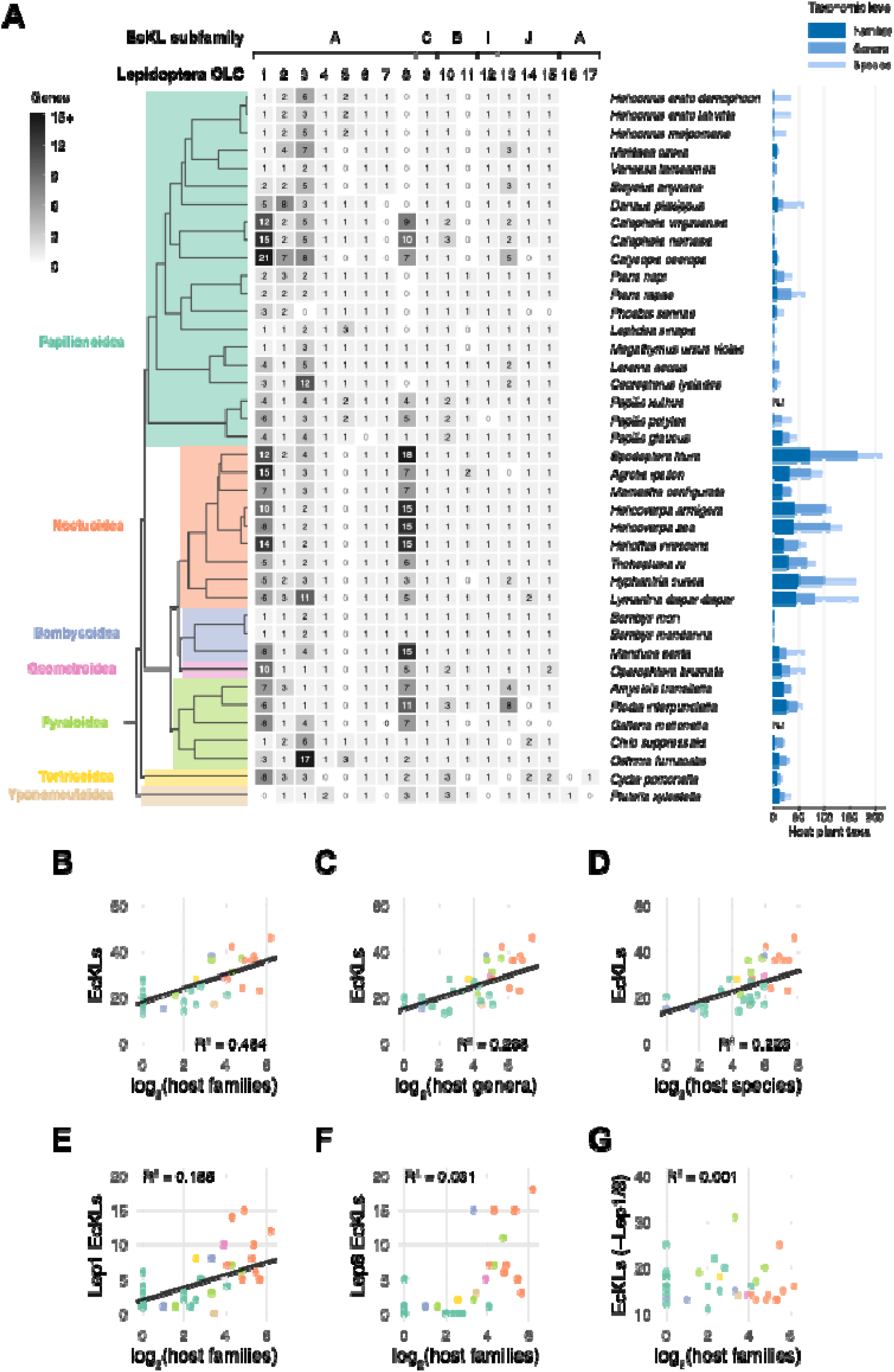
EcKL gene family size correlates with host plant taxonomic diversity in phytophagous Lepidoptera. **A.** (Left) Lepidoptera OLCs for 40 species across seven taxonomic superfamilies. Boxes show the number of genes per OLC per species (indicated by the shading intensity and the number inside). EcKL subfamily groupings per OLC are shown at top. See Fig. S3 for a tree of Lepidoptera OLCs. The phylogenetic tree is derived from Fig. S6; coloured boxes indicate the superfamilies. (Right) The number of host plant families, genera and species for each lepidopteran species, according to the HOSTS database (Robinson et al. 2010). As families ≤ genera ≤ species, the data are displayed as overlayed bars, such that the right-most end of the bar indicates the total count, not the apparent length. ND, no data. **B–G.** PGLS regressions between EcKL gene number and log_2_-transformed counts of host plant families, genera or species. Dots (insect species) are coloured by superfamily as in A. Only significant models (p < 0.05) have their trendline displayed (black lines); for full model outputs, see Table S5. **B–D.** Regressions between total EcKL gene number and transformed host plant families (B), genera (C) or species (D). **E–F.** Regressions between EcKL gene number in the Lep1 (E) or Lep8 (F) OLCs and transformed host plant families. **G.** Regression between total EcKL gene number without Lep1 or Lep8 genes and transformed host plant families.

For our full dataset of 38 lepidopteran species, EcKL family size was only marginally significantly associated with genera (R^2^ = 0.099, p = 0.031) but not species (R^2^ = 0.052, p = 0.052) or families (R^2^ = 0.06, p = 0.075). After further investigation, we realised three outlier taxa—*Calephelis nemesis*, *Calephelis virginiens* and *Calycopis cecrops* —were likely not purely herbivorous but also detritivorous (Cong et al. 2016; Kendall 1959) and could be confounding the analysis. When fitting similar models excluding these three species (n = 35), EcKL family size was significantly positively associated with host families (R^2^ = 0.484, p = 2.1×10^-6^; Fig. 5B), genera (R^2^ = 0.288, p = 5.3×10^-4^; Fig. 5C) and species (R^2^ = 0.223, p = 2.5×10^-3^; Fig. 5D).

We also fit models to explain the size of each of seven Lepidoptera OLCs (Lep1, Lep2, Lep3, Lep5, Lep8, Lep10 and Lep13) that show moderate to high copy number variation across species (at least three genes in at least one species; Fig. 5A) using log_2_-transformed host family count, to explore if any specific clades explain the association in total gene number (Table S5). Only the size of the Lep1 clade was significantly positively associated with host families (R^2^ = 0.185, p = 5.8×10^-3^; Fig. 5E). While the Lep8 clade was not significantly associated with host families (R^2^ = 0.031, p = 0.168; Fig. 5E), further investigation showed it was marginally significantly associated with host genera (R^2^ = 0.092, p = 0.043) and species (R^2^ = 0.102, p = 0.035) and was also strongly positively associated with the size of the Lep1 clade (R^2^ = 0.601, p = 2.7×10^-8^ and R^2^ = 0.396, p = 5.8×10^-3^, depending on variable order). Furthermore, removal of both Lep1 and Lep8 gene count was necessary for total EcKL gene count to no longer be predicted by host families (R^2^ = 0.001, p = 0.318; Table S5), suggesting both clades contribute to the overall positive association. Taken together, these data suggest the size of both the Lep1 and Lep8 clades may have evolved in concert with increased host plant diversity and consequent increases in dietary chemical complexity. Curiously, the Lep1 clade contains *BmEc22K*, the ecdysteroid 22-kinase in *Bombyx mori*.

### EcKLs have low rates of duplication in taxa with chemically stable diets

As a final test of EcKL association with detoxification, we explored the copy number stability of EcKL genes in two well-sampled taxa with chemically stable diets—tsetse flies and bees—and compared them to 12 species in the saprophagous genus *Drosophila* (Markow & O’Grady 2007), the EcKL stability of which has been previously studied (Scanlan et al. 2020). Tsetse flies (genus *Glossina*) are obligate haematophages with adenotrophic viviparity, meaning all dietary compounds originally derive from mammalian blood (Tobe 1978). Apoidea is a hymenopteran lineage that contains bees (Anthophila) and the predatory sphecoid wasps (Sann et al. 2018); the vast majority of bees feed on pollen and nectar, a chemically stable diet relatively low in phytotoxins (Rivest & Forrest 2020), and have relatively few other detoxification genes (Claudianos et al. 2006; Berenbaum & Johnson 2015; Darragh et al. 2021). Consistent with limited XD in their ecological niche, we found both tsetse flies (n = 6 species) and bees (n = 14 species) have very few EcKLs relative to their insect order (26–28 and 12–26, respectively). We hypothesised that these two lineages should also show very limited signs of EcKL copy number variation (gene gain or loss within ancestral clades) compared with *Drosophila*, due to their diets’ chemical stability.

A phylogenetic analysis of EcKLs in six genomically annotated *Glossina* species inferred 30 GLCs (belonging to 14 dipteran OLCs; Fig. 3A), of which eight have experienced a change in copy number (Fig. S10); overall, one duplication and eight loss events were inferred.

A phylogenetic analysis of EcKLs in 14 genomically annotated bee species placed them into 12 hymenopteran OLCs, of which only two have experienced a change in copy number (Fig. S5); overall, seven duplication and zero loss events were inferred. Due to strong transcriptomic sampling of Apoidea in previous studies (Sann et al. 2018; Peters et al. 2017), we used this data to annotate EcKLs in the TSAs of an additional 75 species, including 39 bees and 36 sphecoid wasp relatives (Fig. S11). Gene absences were confined to transcriptome data only, consistent with absences due to transcript abundance rather than true genomic gene loss, and gene copy number increases were only found in 16 additional instances: these data are consistent with low rates of duplication and loss, although they cannot be used to estimate clade stability measures due to the unknown false negative rate. Interestingly, Hym22 genes (subfamily A) were found only in Apoidea species outside of the clade of Anthophila+Pemphredonidae+Philanthidae (Fig. S11), consistent with loss of Hym22 before the most recent common ancestor of bees. We used the inferred gene duplication and loss events in genomically annotated species in each of *Glossina*, Anthophila and *Drosophila*, as well as the relevant branch lengths from our dated phylogeny (67, 788 and 363 m.y., respectively; Fig. S6), to estimate rates of duplication and loss per clade per million years (m.y.). *Glossina* experienced 5.0×10^-4^ duplications/clade/m.y. and 4.0×10^-3^ losses/clade/m.y.; Anthophila experienced 7.4×10^-4^ duplications/clade/m.y. and 0 losses/clade/m.y.; and *Drosophila* experienced 4.0×10^-3^ duplications/clade/m.y. and 4.3×10^-3^ losses/clade/m.y. Duplication rates in the chemically stable taxa were therefore 12.5% and 18% that of *Drosophila*, while loss rates were 93% and 0% that of *Drosophila*. This low duplication rate across both taxa is consistent with our hypothesis.

## Discussion

Understanding and contextualising gene and protein function in a large gene family requires a working knowledge of its evolutionary history. In this study, we have performed the first comprehensive phylogenomic analysis of the EcKL gene family across insects. The classification of EcKLs into subfamilies—whose retention and copy number conservation vary dramatically within and between insect orders— raises some functional hypotheses. The high single-copy retention of subfamily I and subfamily C EcKLs across many insect taxa suggests they could have developmentally or physiologically important endogenous substrates, similar to the Halloween P450s that synthesise ecdysteroids from dietary sterols (Rewitz et al. 2007). Notably, no EcKL subfamilies appear perfectly conserved across all insects. One explanation is that these gene absences are due to genome assembly errors and some EcKLs are indeed present in all insect genomes. This is unlikely to be a complete explanation, given the lack of subfamily I in brachyceran dipterans includes *Drosophila melanogaster*, which has the most completely sequenced and well-analysed genome of all insects. The more likely explanation is that no EcKL has a function that is essential in all insect taxa—indeed, the high variability in subfamily size between various insect orders suggests different subfamilies may have important lineage-specific functions that have resulted in substantial ‘gene blooms’, a phenomenon also seen in the P450s (Feyereisen 2011) .

We have also identified EcKL clades that are conserved as single-copy orthologs within (but not necessarily between) the insect orders of Diptera, Lepidoptera and Hymenoptera. We hypotheses these genes are involved in important, order-specific developmental or metabolic processes. One example is Dip12 (subfamily B) in Diptera, represented by *CG2004* in *D. melanogaster*, which may modulate the endoplasmic reticulum stress pathway through protein-protein interactions (Chow et al. 2016, 2013), but *CG2004*’s substrate-level function has not been characterised. This variability in both total EcKL number and the size of specific subfamilies— subfamilies A and H have some of the largest differences between orders—requires an explanation. This study provides phylogenetic comparative evidence that EcKL gene family size is positively associated with the assumed xenobiotic diversity (XD) of an insect’s ecological niche. We found evidence that the size of the GSTs, and to a lesser extent the P450s, evolves in response to a need for greater XD. Previous studies have found some associations between insect diet and detoxification gene family sizes, but did not do so with phylogenetically corrected statistical methods as we performed here size (Rane et al. 2019; Breeschoten et al. 2021; Calla et al. 2017). Our findings should be further studied with better measures of XD, such as direct measurements of xenobiotic exposure across a range of taxa. Specifically, our use of the HOSTS database (Robinson et al. 2010) for lepidopteran species could be expanded to species with new high-quality genome assemblies and could search for specific clades within the classical detoxification gene families that are associated with host plant diversity.

The poor association of P450 family size with XD found in this study is surprising, as P450s are the most well-characterised detoxification genes in eukaryotes. We might explain this finding in a few ways: (1) our estimation of XD was inaccurate and does not reflect the true XD of an organism in nature; (2) P450s are involved in other lineage-specific metabolic processes (eg. Beran et al. 2019; Fu et al. 2019) that produce changes in copy number that obscured the detoxification signal; and/or (3) other factors, such as microbiome-based xenobiotic metabolism (eg. Ceja-Navarro et al. 2015; Motta et al. 2022) or ‘social detoxification’ mechanisms (Berenbaum & Johnson 2015), reduce the need for a large P450 family in some insects. Given we observed robust associations between diet/XD and EcKL and GST gene family sizes, explanation 2 may be the most likely.

Curiously, we found no evidence that CCE gene number changes with diet or XD across all insects, casting doubt on their general role as detoxification enzymes for ecologically relevant toxins, despite their clear involvement in the evolution of resistance to some synthetic insecticides (Oakeshott et al. 1999, 2005). However, two recent studies have independently found a positive association between polyphagy and CCE family size in Lepidoptera (Dort et al. 2023; Breeschoten et al. 2021), which might suggest detoxification roles for CCEs are limited to specific insect taxa.

The robust positive associations between EcKL gene family size and diet/XD, as well as lower estimated EcKL duplication rates in taxa with chemically stable, compared with unstable, diets, support our hypothesis that EcKLs are involved in detoxification processes in insects. This hypothesis was previously motivated and supported by data in *Drosophila* (Scanlan et al. 2020, 2022) and strongly suggests that phosphorylated metabolites of xenobiotic compounds found across insects are both produced by EcKL enzymes and also contribute to metabolic detoxification in natural contexts. Despite this, EcKL enzymes with xenobiotic substrates are yet to be biochemically characterised. The current work provides strong phylogenetic motivation to functionally characterise various EcKLs for detoxification roles, with the most striking candidates belonging to the Dip1/3/4/7 clades in Diptera (subfamily H) and the Lep1/8 clades in Lepidoptera (subfamily A). The former are closely related to EcKLs that mediate caffeine tolerance in *D. melanogaster* (Scanlan et al. 2022) and show substantial changes in copy number between dipteran species. The latter are positively associated with host plant diversity and likely contribute to phytochemical detoxification in many polyphagous species of Lepidoptera. Conspicuous expansions of subfamily A and/or H EcKLs in cockroaches, fungus-cultivating ants, sawflies, beetles and stick insects are also worthy of further study. Additionally, the recent discovery that the whitefly *Bemisia tabaci* (order Hemiptera) detoxifies cyanogenic glucosides through phosphorylation (Easson et al. 2021) should prompt the study of specific EcKL enzymes in this process.

Despite support for the detoxification hypothesis here and in previous studies (Scanlan et al. 2020, 2022), the best biochemically established function for EcKL enzymes remains ecdysteroid kinase activity. Two genes, *BmEc22K* and *AgEcK2*, are known to encode enzymes that phosphorylate ecdysteroids at the C-22 position (Peng et al. 2022; Sonobe et al. 2006). Our phylogenetic analyses placed these genes in different EcKL subfamilies—subfamily A (*BmEc22K*) and subfamily D (*AgEcK2*)—which suggests other genes in these two subfamilies are likely to encode ecdysteroid kinases. It also demonstrates very similar enzyme functions can be found in different clades of the family, either indicating convergent evolution of substrate specificity or conservation of the function of their common ancestor; the latter would imply the ancestral function of subfamilies B, C, L and M is also ecdysteroid kinase activity. Peng et al. (2022) also identified the presence of 3-dehydro-20E 22-phosphate in *A. gambiae*, which is not a product of AgEcK2, indicating an uncharacterised enzyme is responsible; due to the single-copy conservation of subfamily D in dipterans, the encoding gene must belong to a different EcKL subfamily.

*D. melanogaster* is the insect with the most powerful genetic toolkit and is well positioned to identify additional ecdysteroid kinases in the EcKL family. Four ecdysteroid-phosphate conjugates have been determined or suggested to exist in *D. melanogaster*: E 22-phosphate in adult ovaries (Grau et al. 1995; Pis et al. 1995); 20,26-dihydroxy-E 26-phosphate in the S2 cell line (Guittard et al. 2011); 3-dehydro-E 2-phosphate in larval homogenate (Hilton 2004); and 3-epi-20E 3-phosphate in larvae (Sommé-Martin et al. 1988). *CG5644* is the direct ortholog of *AgEcK2*, making it an excellent ecdysteroid 22-kinase candidate, and while it has very low expression in the ovary as a whole (Leader et al. 2018), it appears enriched in the migratory border cells of the ovary (Borghese et al. 2006; NCBI GEO accession GSE4235), raising the possibility it produces ovarian E 22-phosphate. Intriguingly, the subfamily C EcKL *CG14314* encodes a protein that might physically interact with the ecdysteroid-phosphate phosphatase EPP/CG13604 (Murali et al. 2011; Davies et al. 2007), consistent with a function in ecdysteroid metabolism. Both *CG5644* and *CG14314* are strongly enriched for expression in the adult nervous system (Leader et al. 2018) and might modulate steroidal control of sleep, behaviour and memory (Li et al. 2023; Okamoto & Yamanaka 2020; Ishimoto et al. 2012, 2009). *CG13813* and *Pinkman* are *D. melanogaster* subfamily A genes that could also encode ecdysteroid kinases; notably, *CG13813* is strongly induced by 20E in cell culture (Stoiber et al. 2016; Gauhar et al. 2009) and is enriched for expression in the 3^rd^-instar larval ring gland (Ou et al. 2016), both consistent with a function in ecdysteroid metabolism. Ecdysteroid-phosphate conjugates have also been identified in other insects: the lepidopterans *Bombyx mori*, *Manduca sexta* and *Pieris brassicae*, and the orthopterans *Locusta migratoria* and *Schistocerca gregaria*. Lepidopteran conjugates include the 22-phosphates produced by BmEc22K in *B. mori* (Sonobe et al. 2006), but also 2-, 3- and 26-phosphates (Beydon et al. 1987; Lozano et al. 1989; Thompson et al. 1988; Sonobe & Yamada 2004). While Lep1 orthologs of *BmEc22K* might be responsible for these additional conjugates, genes in other subfamily A clades (Lep2 through Lep8) are also good candidates. Orthopteran phosphate conjugates also include 2-, 3- and 22-phosphates (Gibson et al. 1984; Modde et al. 1984; Isaac et al. 1982, 1983; Isaac & Rees 1985). Our analysis of transcriptomic data from Orthoptera suggests that this taxon might not possess subfamily A, meaning these phosphate conjugates must be produced by enzymes in other EcKL subfamilies.

Our finding that EcKLs are restricted to the Tetraconata (Hexapoda + Crustacea) is consistent with the known distribution of ecdysteroid-phosphates conjugates in arthropods—ie. present in insects and crustaceans (Subramoniam 2000; Young et al. 1991, 1993; Suzuki et al. 1996) but (to our knowledge) as-yet-undetected in myriapods or chelicerates. Further phylogenetic and functional analyses are required to explore whether ecdysteroid kinase activity is the ancestral enzymatic function of the EcKLs, and whether detoxification functions have also evolved for non-insect EcKLs.

A surprising finding in this study is the strong association between host plant diversity and the size of the Lep1 EcKL clade (subfamily A) in Lepidoptera, which suggests this clade has expanded to mediate phytochemical chemical detoxification in multiple species, despite containing a biochemically characterised ecdysteroid 22-kinase, BmEc22K. Some plants are known to biosynthesise phytoecdysteroids, which act as ecdysteroid agonists and disrupt insect herbivore growth (Adler & Grebenok 1995; Dinan 2001). This may have produced an evolutionary bridge through which Lep1 paralog expression and substrate specificity could shift to digestive tissues and xenobiotic compounds, respectively. Ingested ecdysteroids are primarily phosphorylated in the midguts of some (Webb et al. 1996, 1995; Weirich et al. 1986; Williams et al. 1997)—but not all (Rharrabe et al. 2007)—lepidopterans. Notably, however, noctuid moths are highly resistant to phytoecdysteroid ingestion through the formation of fatty acyl conjugates, not phosphates (Duan et al. 2020; Robinson et al. 1987). The large number (8–14) of Lep1 EcKLs in this insect clade therefore suggests these enzymes have other phytochemical substrates. A genomic locus containing four Lep1 EcKLs in the noctuid *Helicoverpa armigera*, subspecies *H. armigera conferta* has recently undergone a selective sweep (Anderson et al. 2018), raising the possibility of metabolic adaptation to a novel diet; alternatively, this locus may have been selected for a role in hormone metabolism.

In this work, we have conducted the first comprehensive phylogenomic analysis of the EcKL gene family and provided additional support for the hypothesis that EcKLs encode detoxicative kinases. To build on these results, focused studies should be undertaken to identify the biochemical substrates of candidate detoxification EcKLs, particularly in Lepidoptera, where strong associations have been found between dietary phytochemical diversity and EcKL copy number. Future studies could also expand on the framework established here and explore the co-evolution of insect and plant genomes and metabolomes using comparative phylogenetic methods; such studies might uncover additional, previously undetected enzyme families associated with xenobiotic metabolism and insect-plant interactions.

## Materials and Methods

### Genome and transcriptome annotation

Arthropod genome assemblies uploaded to NCBI Assembly (Kitts et al. 2016), VectorBase (Giraldo-Calderón et al. 2015) or Lepbase (Challis et al. 2016) were queried with EcKL protein sequences using tblastn (Altschul et al. 1990) and matching scaffolds/contigs were downloaded and annotated manually in Artemis (Carver et al. 2012). RefSeq proteins and/or transcriptome shotgun assemblies (TSAs) from the closest available taxonomic group were queried with rough annotated gene model translations to iteratively inform intron-exon boundaries. Gene models were classified as ‘full’, ‘partial’ or ‘pseudogenous,’ with the latter defined as containing two or more inactivating mutation (frameshift, splice donor/acceptor loss, premature stop codon etc.); pseudogenous models were not rigorously annotated or collated. Gene models with only a single inactivating mutation were considered null alleles (or potentially sequencing errors), had the mutation conservatively reverted, and otherwise treated as ‘full’ models. Only full or partial gene models were counted towards EcKL totals for each species. Rarely, there was evidence of alternatively spliced gene models that affected the protein sequence, in which case isoforms were counted as individual genes towards gene totals if alternative protein-coding exons accounted for more than 25% of the coding sequence. For genome assembly information, including the total number of full and partial gene models per assembly, see Table S1.

To obtain EcKL subfamily sequences from insect orders not represented in the genome-annotation dataset, NCBI TSAs were tblastn queried with putative EcKL subfamily sequences from closely related orders, and the putative open reading frames from selected transcripts were translated with the ExPASy Translate tool (Artimo et al. 2012)—up to three sequences with the highest sequence similarity per subfamily were collated per order.

The presence or absence of EcKLs within the taxonomic classes of Arthropoda was determined by using tblastn against TSAs from each class, using the amino acid sequences of *CG5644* from *Drosophila melanogaster* (subfamily D) and *GB49960* from *Apis mellifera* (subfamily I) as queries. The top hits were used as BLAST queries against the Arthropoda nr database to check for RNA contamination from the diet; E-values below 0.05 were considered significant. TSAs of taxonomic classes were accessed in NCBI on 2023-03-15.

Apoidea transcriptomes were annotated by using *Apis mellifera* EcKLs as tblastn queries against assembled transcriptomic contigs in the NCBI database and translating open reading frames with the ExPASy Translate tool. Overlapping partial transcripts were manually merged into larger contigs where possible. EcKLs from transcriptomes were assigned to ancestral hymenopteran clades by aligning with *A. mellifera*, *Nasonia vitripennis* and *Neodiprion lecontei* EcKLs with MAFFT (Katoh & Standley 2013) and constructing phylogenetic trees with IQ-TREE (Nguyen et al. 2015), ala. the methods described below.

### Multiple sequence alignment, phylogenetic inference and ancestral clade assignment

Before alignment, sequences with multiple EcKL domains were split into smaller sequences each containing one EcKL domain, and N-or C-terminal regions that did not share homology with the rest of the EcKL family were removed. Sequences were aligned with MAFFT 7.402, using the L-INS-i setting unless otherwise specified (recommended for sequences with a single conserved domain; Katoh & Standley 2013). MSAs were manually trimmed in AliView (Larsson 2014) to remove obviously poorly aligned columns (typically at the N-and C-terminal ends of the alignment), but over-trimming (removal of >20% of columns) was avoided, given that can reduce the accuracy of phylogenetic inference (Dessimoz & Gil 2010; Tan et al. 2015). Phylogenetic inference was performed on trimmed MSAs with IQ-TREE 1.6.10 (Nguyen et al. 2015), using the ModelFinder program to find the best model for each MSA (Kalyaanamoorthy et al. 2017) and UFBoot2 for bootstrapped branch support values (Hoang et al. 2018). IQ-TREE runs were performed 5–10 times per MSA and the tree with the highest log-likelihood was used. MAFFT and IQ-TREE runs were performed in the CIPRES Science Gateway computing environment on the XSEDE platform (Miller et al. 2010).

For ancestral clade phylogenetics where the total number of sequences was approx. >500, all sequences in the taxon were aligned, then the MSA was pared down to a subset of representative sequences (ie. groups of sequences with very high similarity were reduced to a single representative, and partial sequences were also removed) in order to improve phylogenetic parameter estimation (recommended by Burnham & Anderson 2004). Clade designations for excluded sequences (ie. those not in the subset) were determined by their grouping with similar sequences in the guide tree produced by MAFFT during alignment. Where ambiguities arose, small groups of sequences from the total sequence set (eg. all sequences thought to be in a particular clade, plus a small number of appropriate outgroup sequences) were aligned and a tree was produced with IQ-TREE.

To determine insect EcKL subfamilies, a rough initial tree was constructed of a single sequence from each ancestral clade from Diptera, Lepidoptera and Hymenoptera, as well as representative sequences (ala. ancestral clade phylogenetics above) from all other genome-annotated insect taxa. To produce the final EcKL subfamily tree seen in Fig. 2A, a single sequence from each putative subfamily per order (including transcriptome-only orders) was selected, except for Lepidoptera, where a single representative sequence from each of the Lep1–8 and Lep6–17 ancestral clades (subfamily A) was included in order to break up long branch lengths associated with these sequences, and for Odonata, Ephemeroptera, Zygentoma and Archaeognatha, where up to three sequences were selected from these orders per subfamily to help break up long branches. In total, 241 sequences were selected for this ‘core set’ (Supplementary Data) and aligned using the E-INS-i setting (for better deep node support; Wilding et al. 2017) in MAFFT, and then imported into IQ-TREE for analysis. To classify individual EcKLs into subfamilies, groups of sequences from each order were independently aligned to the ‘core set’ and a tree produced with IQ-TREE.

Pairwise amino acid identity of EcKL proteins between and within subfamilies was calculated using the full-length sequences in the ‘core set’ described above (n = 202 sequences) and the ‘seqidentity’ function in the *bio3d* package (version 2.4-4) in R (Grant et al. 2006).

### An estimated time tree for insects

A dated phylogenetic tree with branch lengths in units of millions of years (Fig. S6; Supplementary Data) was manually constructed in Mesquite (Maddison & Maddison 2019) for all 140 insect species annotated for EcKLs, using phylogenetic relationships and node ages published in the literature (Cranston et al. 2011; Espeland et al. 2018; Johnson et al. 2018; Junqueira et al. 2016; Kawahara et al. 2019; Kohli et al. 2016; Krosch et al. 2012; Misof et al. 2014; Moreau et al. 2006; Nygaard et al. 2015; Peters et al. 2017, 2018; Tamura et al. 2004; Wiegmann et al. 2011; Zhang et al. 2018). Where node ages conflicted between studies, the most recent study was used. Nodes with unknown age were estimated based on the known divergence times of similar taxonomic ranks in other parts of the tree (inaccuracy of these node ages is thought to have relatively minor impact on downstream comparative analyses; see Stone 2011).

### Detoxification gene family sizes, dietary groups and XD groups

EcKL gene number per genome (n = 140 species) were determined from genome annotation described above. Cytochrome P450 (P450), glutathione S-transferase (GST) and carboxyl-cholinesterase (CCE) gene family sizes for species with EcKL annotations determined in this study (n = 98 species) were taken from Rane et al. (2019); the UDP-glycosyltransferase (UGT) and ABC transporter (ABC) family sizes from that study were not used, as the data may be inaccurate (Rane et al. 2019). P450, GST and CCE data was also not used for *Gerris buenoi*, as the unrealistically low number of GST genes (six) cast doubt on the reliability of the annotation for this genome.

Dietary groups for each insect were defined as ‘carnivorous’ (carnivorous or parasitoidal; n = 19), ‘herbivorous’ (herbivorous or fungivorous; n = 54), ‘haematophagous’ (n = 9), ‘pollen-feeding’ (n = 15) or ‘detritivorous’ (detritivorous, saprophagous or omnivorous; n = 43; Table S1). If juvenile and adult diets were different, the diet with the higher chemical complexity was used, in the preference order detritivorous > herbivorous > carnivorous > pollen-feeding > haematophagous. *Calephelis nemesis*, *Calephelis virginiens* and *Calycopis cecrops* (Lepidoptera) were coded as detritivorous (Kendall 1959; Cong et al. 2016).

An estimation of ‘xenobiotic diversity’ (XD) was assigned for every species in the dataset, with three levels—small (XD_S_), intermediate (XD_I_) and large (XD_L_)—based on the estimated overall chemical complexity of their diets during juvenile and adult life stages (Table S1). Detritivores and saprophages were coded as XD_L_, except for *Drosophila mojavensis* and *Drosophila sechellia*, which are relatively specialised in their diets compared to related species (Date et al. 2013; Dworkin & Jones 2008) and were coded as XD_I_. Pollen feeders and haematophages were consistently coded with XD_S_. For lepidopteran species, XD was coded based on the host plant diversity (number of host plant families) in the HOSTS database (Robinson et al. 2010): one family, XD_S_; two to nine families, XD_I_; ten or more families, XD_L_. Non-lepidopteran herbivores were typically coded as XDI unless their diets were known to be particularly chemically simple (eg. the wood-eating *Zootermopsis nevadensis*; XD_S_) or complex (eg. the generalist pest *Halyomorpha halys*; XD_L_). Coding XD for some ants and all parasitoid wasps was challenging and the coded values are of low confidence—most ants are omnivorous but can have a preference for carnivory (Clay et al. 2017), while others are technically fungivorous but may be exposed to many different plant secondary metabolites through leaf foraging (De Fine Licht & Boomsma 2010), while the amount of exposure of parasitoids to xenobiotic compounds is poorly understood and is hard to estimate. The uncertainty of these XD assignments was explored during analyses.

### PGLS regression of gene family sizes

Phylogenetic generalised least squares (PGLS) regression between gene family sizes was performed with the ‘pgls’ function in the package *caper* (version 1.0.1) in R, with the λ parameter estimated with maximum likelihood. The *ape* package (v5.0; Paradis & Schliep 2018) was used to manipulate phylogenetic trees in R.

### PGLS regression of gene family sizes and diet/xenobiotic diversity (XD)

Phylogenetic ANOVAs for gene family size against diet or DB were performed with the ‘gls’ function in the *nlme* package (version 3.1-141) in R, with the phylogeny in Fig. S6, pruned to overlap species for P450s, GSTs and CCEs with *ape*, used as a correlation structure with ‘corPagel’ from the *ape* package and a λ value estimated by maximum likelihood. The ‘glht’ function from the *multcomp* package (v1.4-10; Hothorn et al. 2008) in R was used to perform two-sided multiple comparison of means post-hoc tests between each pair of diets or XD levels, with p-values adjusted for multiple comparisons with the ‘single-step’ argument. For XD analyses, two datasets were analysed: the full dataset with all species (140 or 98 species, for EcKLs and P450s/GSTs/CCEs, respectively), and a restricted dataset without omnivorous/fungivorous ants or parasitoid wasps.

### PGLS regression of EcKL family size and host plant diversity in Lepidoptera

Caterpillar and host plant relationships were extracted from the HOSTS database (Robinson et al. 2010) in early 2019, and the number of unique host plant families, genera and species were compiled for each genome-annotated lepidopteran species. Those without an entry in the HOSTS database (*Papilio xuthus*) or no associated plant taxa (*Galleria mellonella*) were not included in further analyses. Detritivory at adult or larval stages may be a confounding variable in this analysis, as rotting food substrates are a likely source of bacterial and fungal toxins, which may influence detoxification gene family evolution. To account for this, we conducted subsequent analyses twice: once on the full dataset, and once on a subset that contained only non-detritivorous species. As related members of the Riodinini tribe are detritivorous as larvae and adults (Hall & Willmott 2000) and species of in the genus *Calephelis* can feed on partially rotten leaves (Kendall 1959), the species *Calephelis nemesis* and *Calephelis virginiens* were marked as detritophagous, as was *Calycopis cecrops*, which feeds on fallen leaves and detritus as larvae (Cong et al. 2016).

Phylogenetic relationships between lepidopteran species were extracted from the master phylogeny previously estimated for all insects (Fig. S6). Host plant families, genera and species counts (‘host counts’) were log_2_-transformed, except where noted. PGLS regression of EcKL number to host counts was performed with the ‘pgls’ function in the package caper, with the λ parameter estimated with maximum likelihood; *ape* was used to manipulate phylogenetic trees.

### Data analysis and visualisation

Data were analysed as described above in the R computing environment (R Core Team 2022), additionally using the *tidyverse* collection (Wickham et al. 2019) and *patchwork* (Pedersen 2022) for data manipulation and visualisation. *ggtree* (v3.6.2; Yu 2020; Yu et al. 2018, 2017) and *treeio* (v1.22.0; Wang et al. 2020) were used to manipulate, visualise and annotate phylogenetic trees.

## Data Availability Statement

The data underlying this article are either available in the article and in its online supplementary material, or will be shared on reasonable request to the corresponding author.

## Acknowledgments

We thank David Heckel and Rebecca Gledhill-Smith for feedback and helpful comments on this project.

