## Supplementary figures and images for "Phylogenomics of the ecdysteroid kinase-like (EcKL) gene family in insects highlights roles in both steroid hormone metabolism and detoxification"

### Figure S1

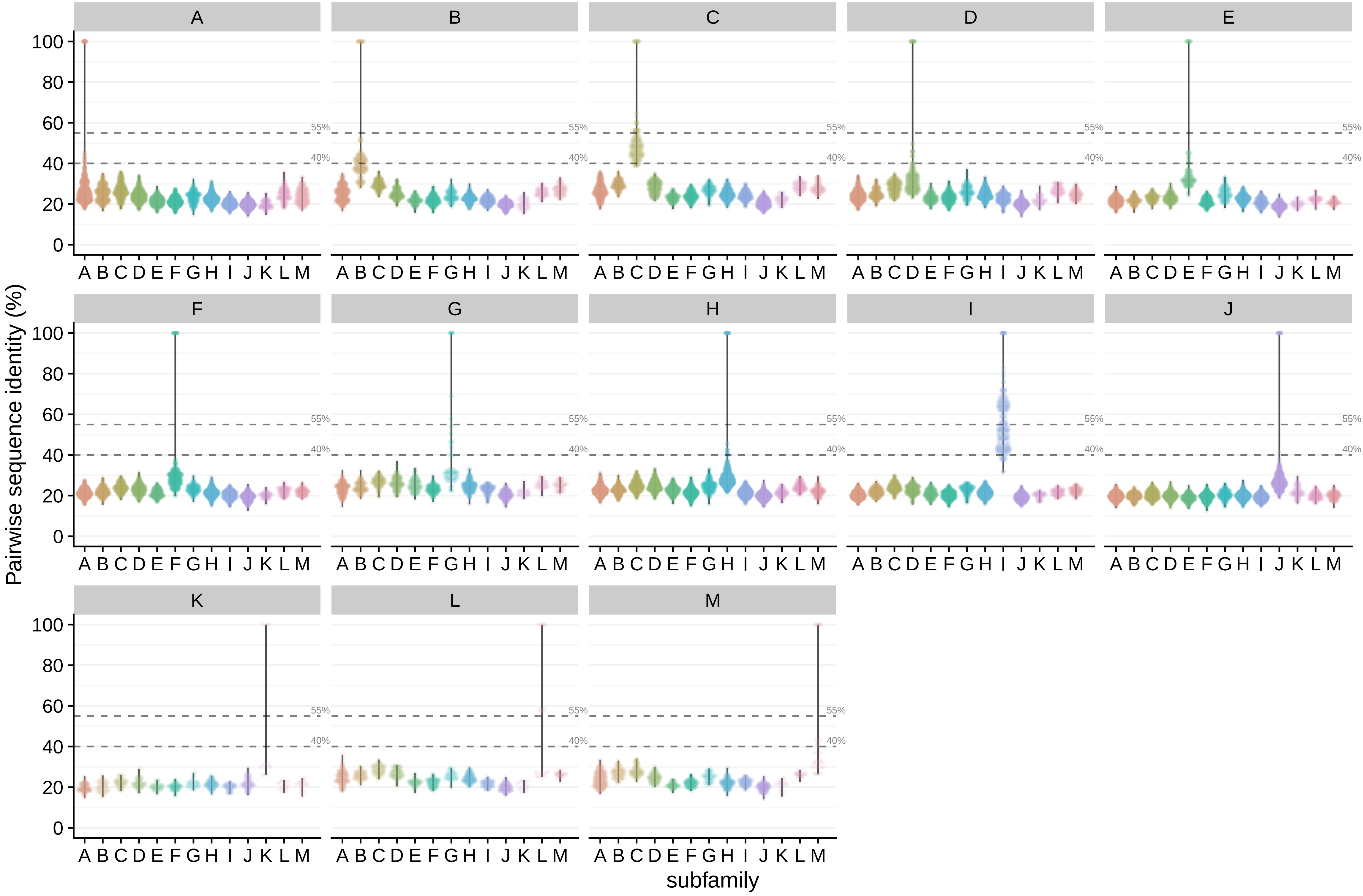

### Figure S2

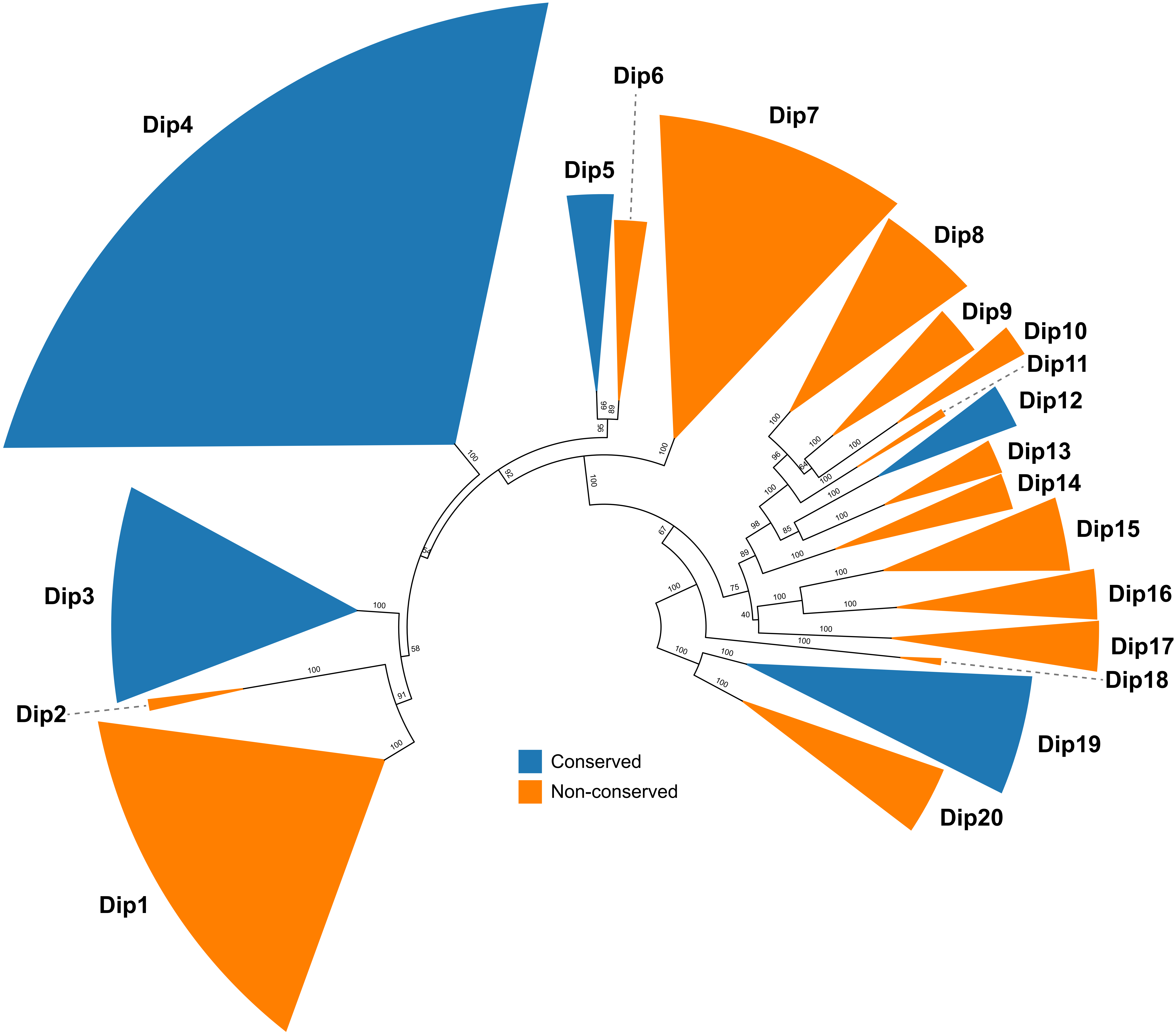

### Figure S3

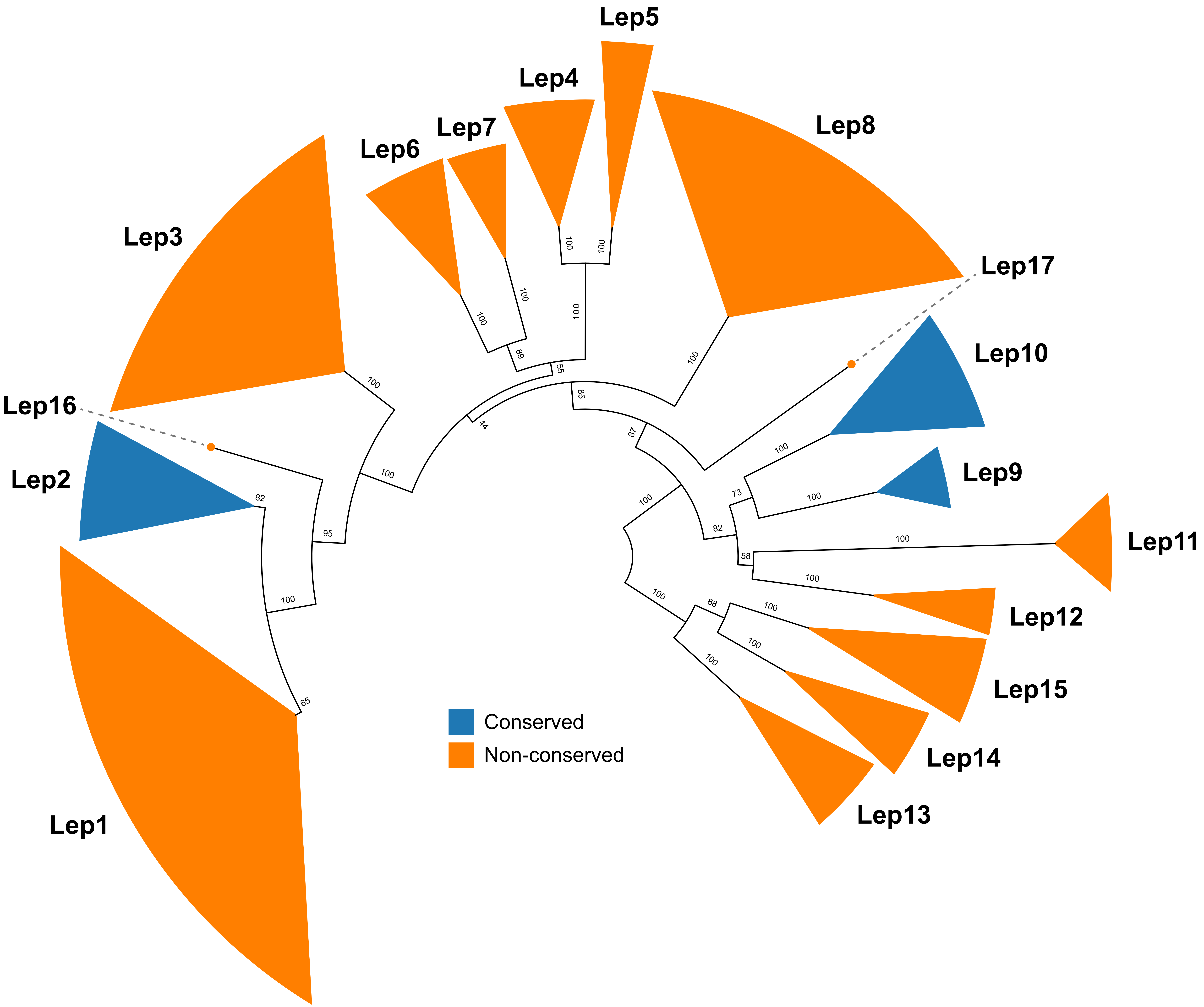

### Figure S4

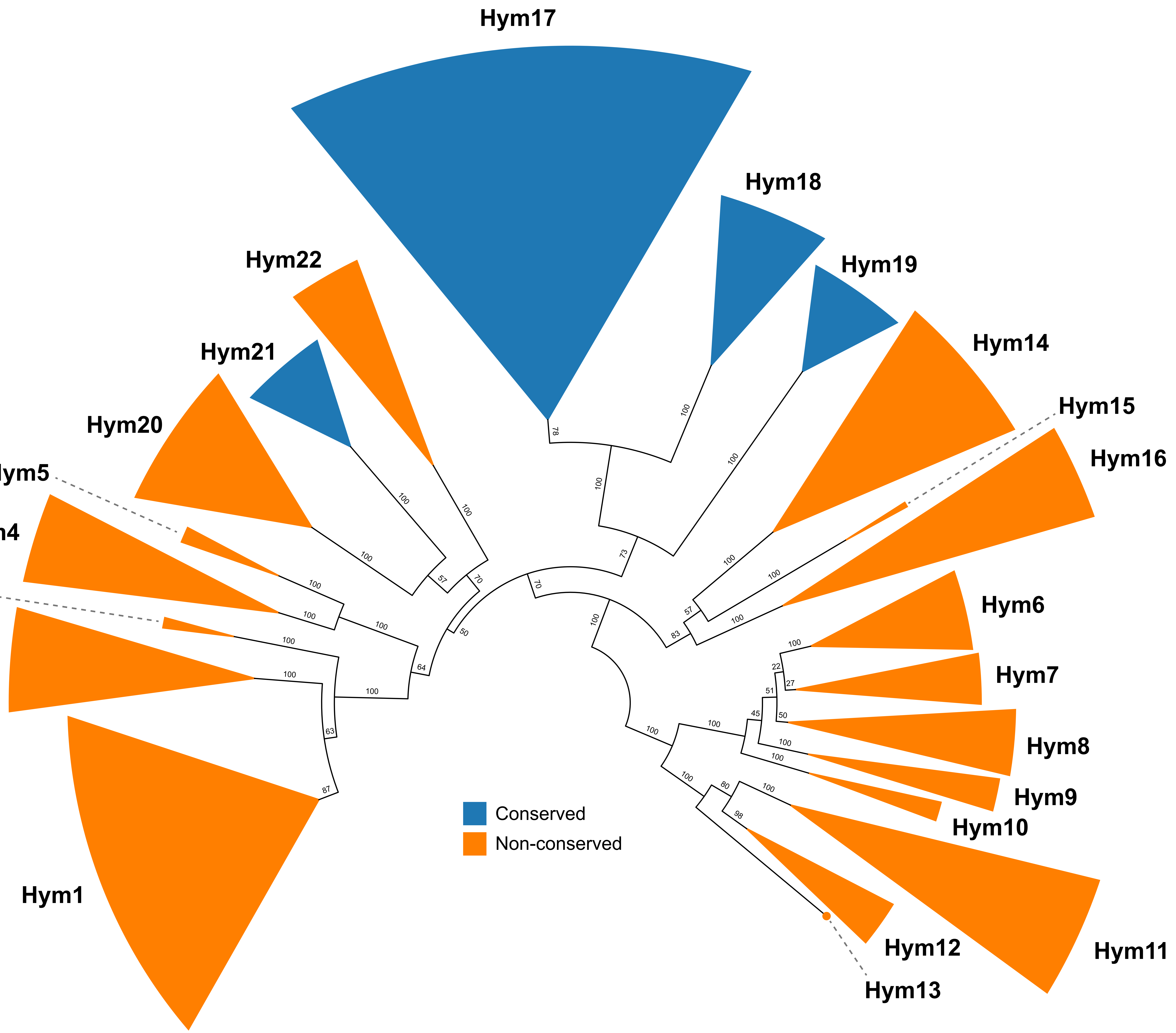

### Figure S5

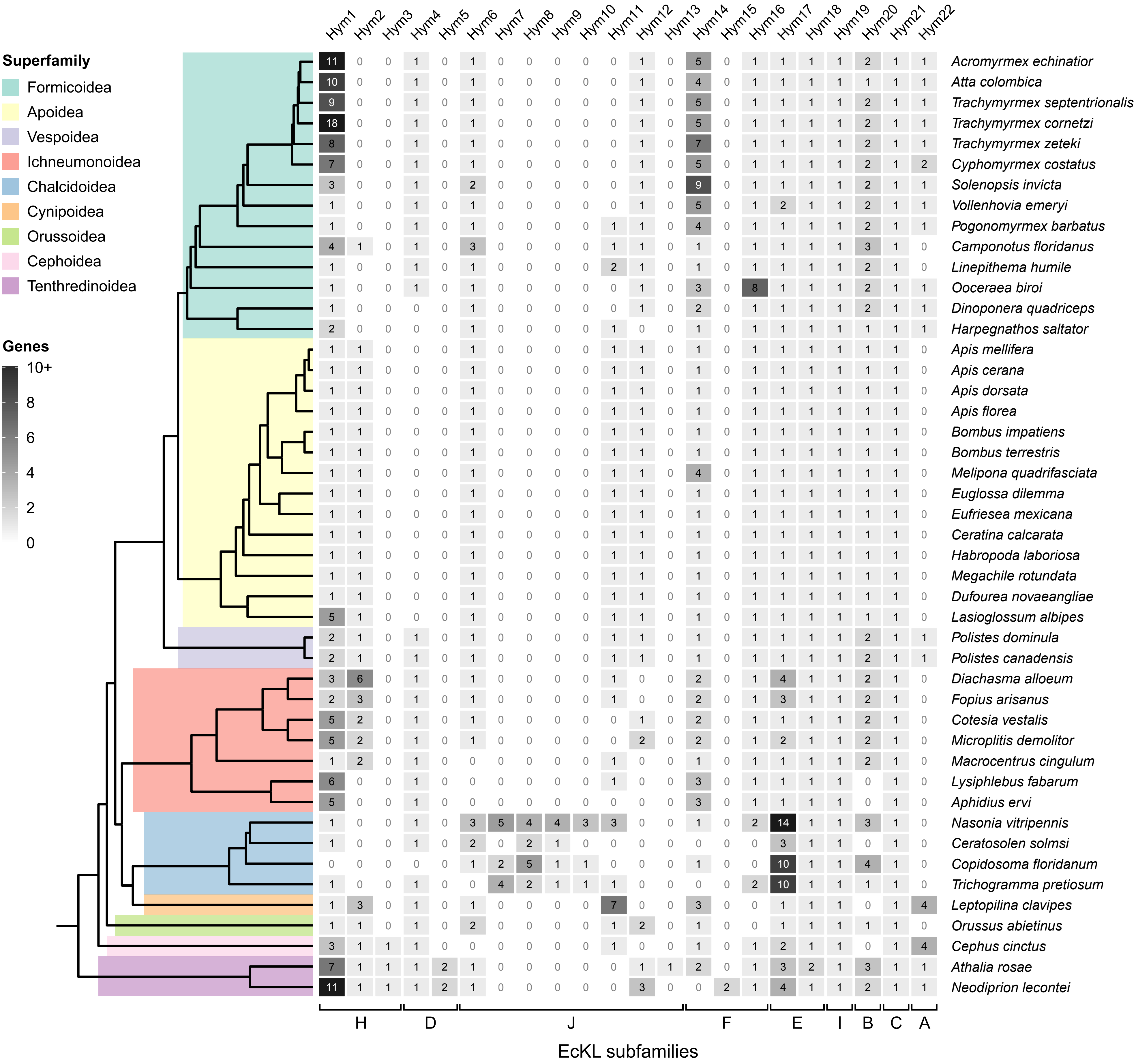

### Figure S6

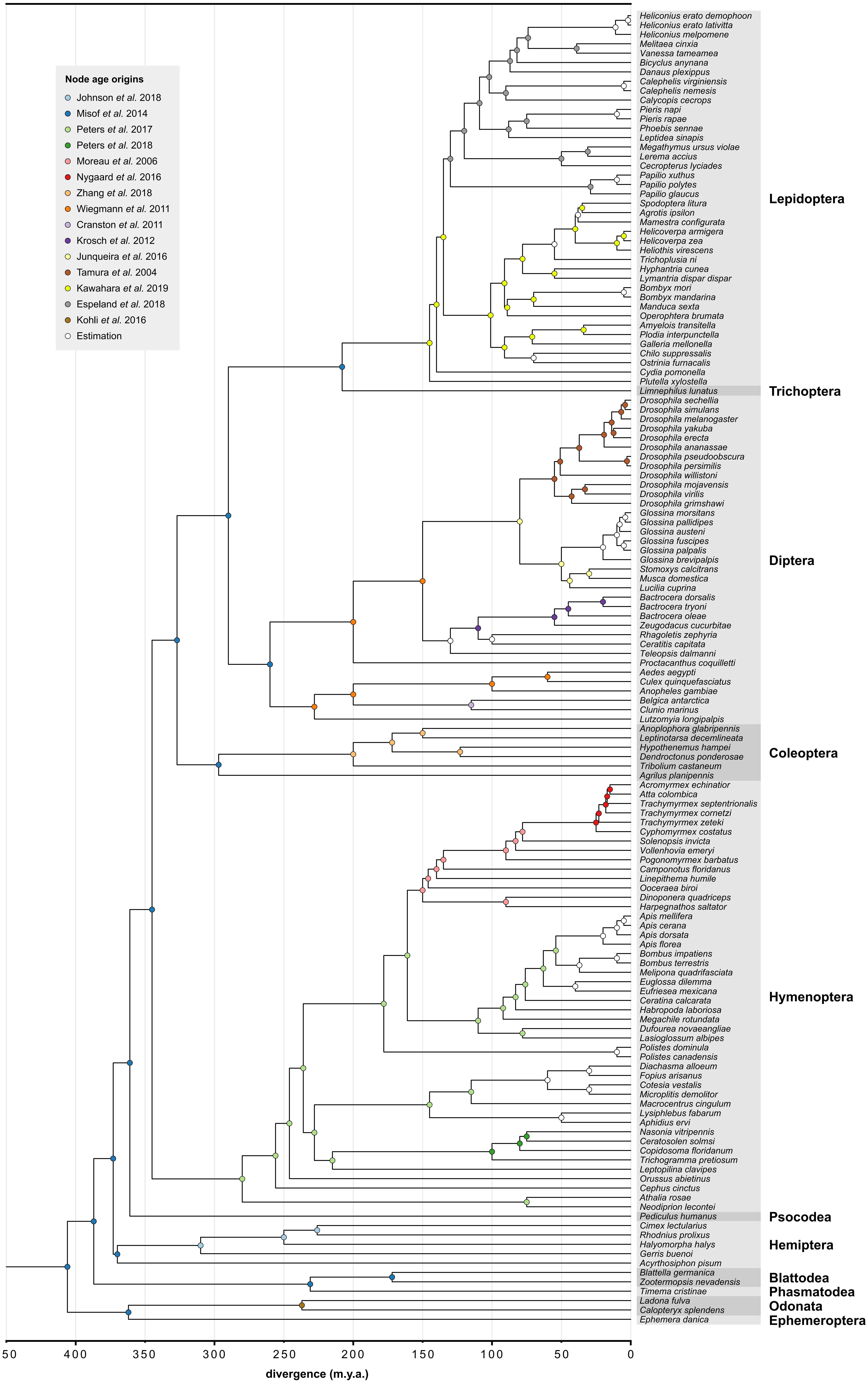

### Figure S7

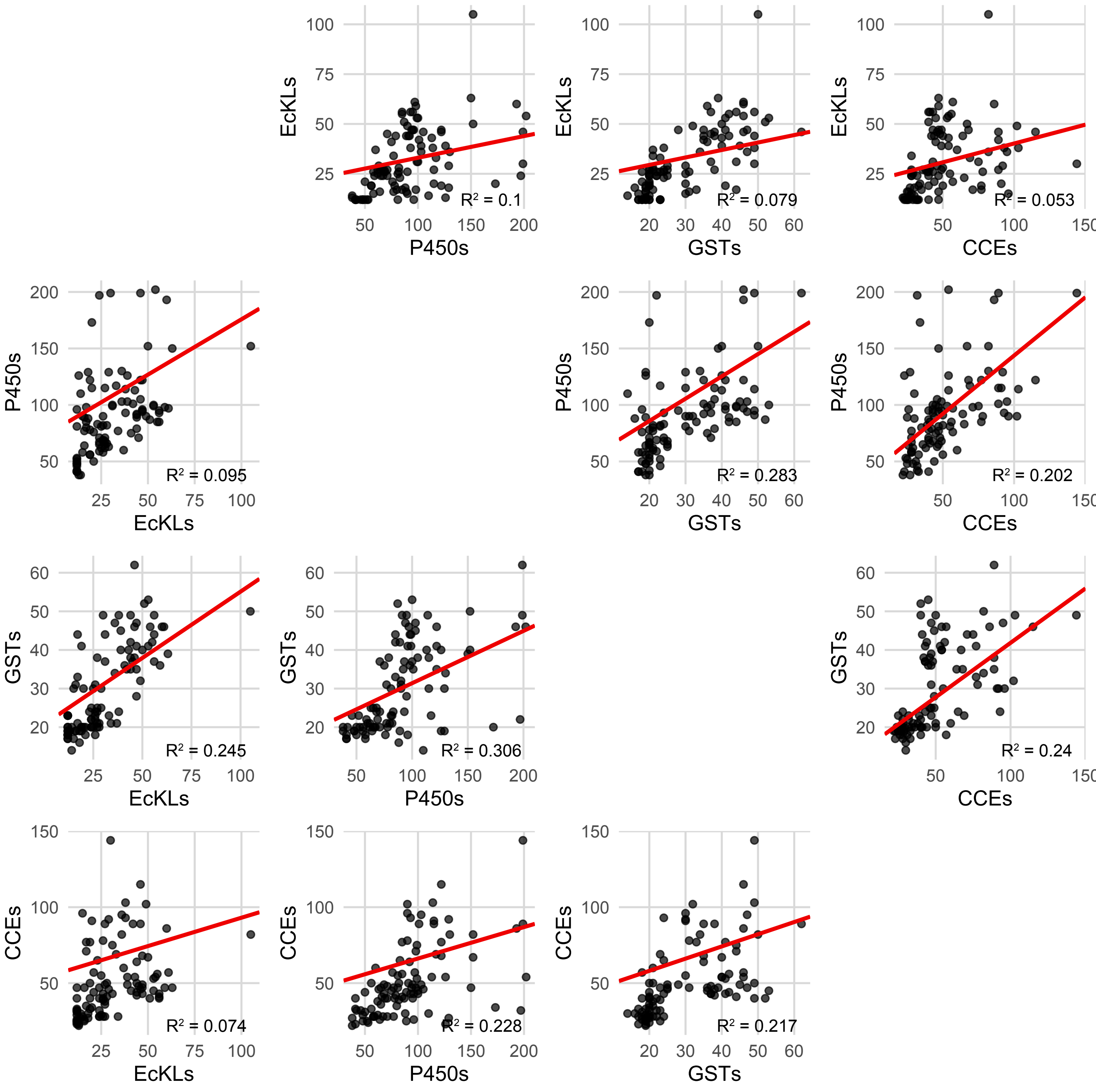

### Figure S8

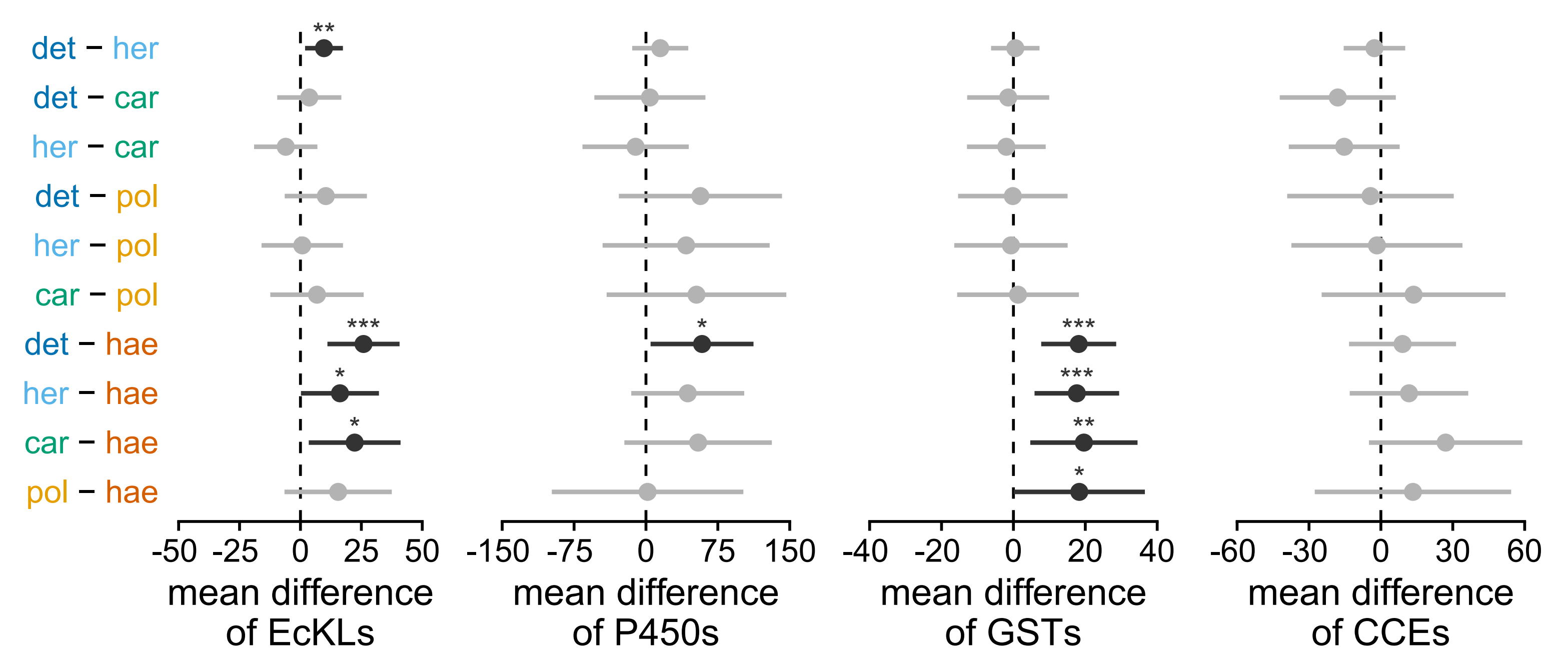

### Figure S9

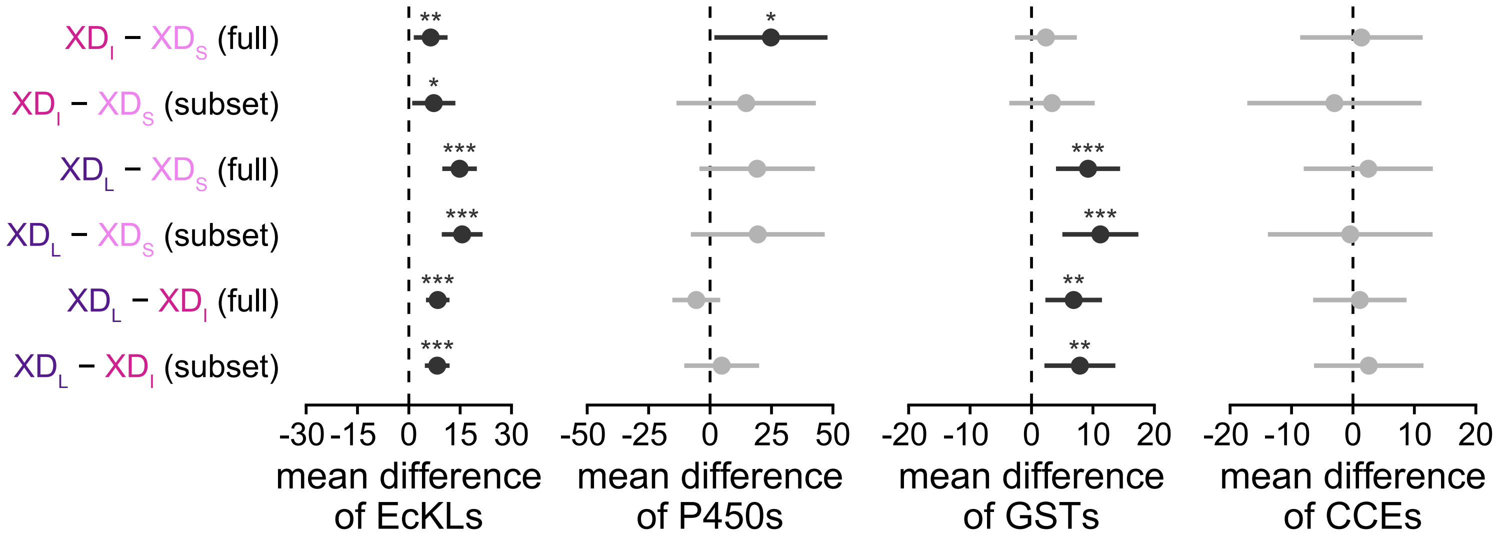

### Figure S10

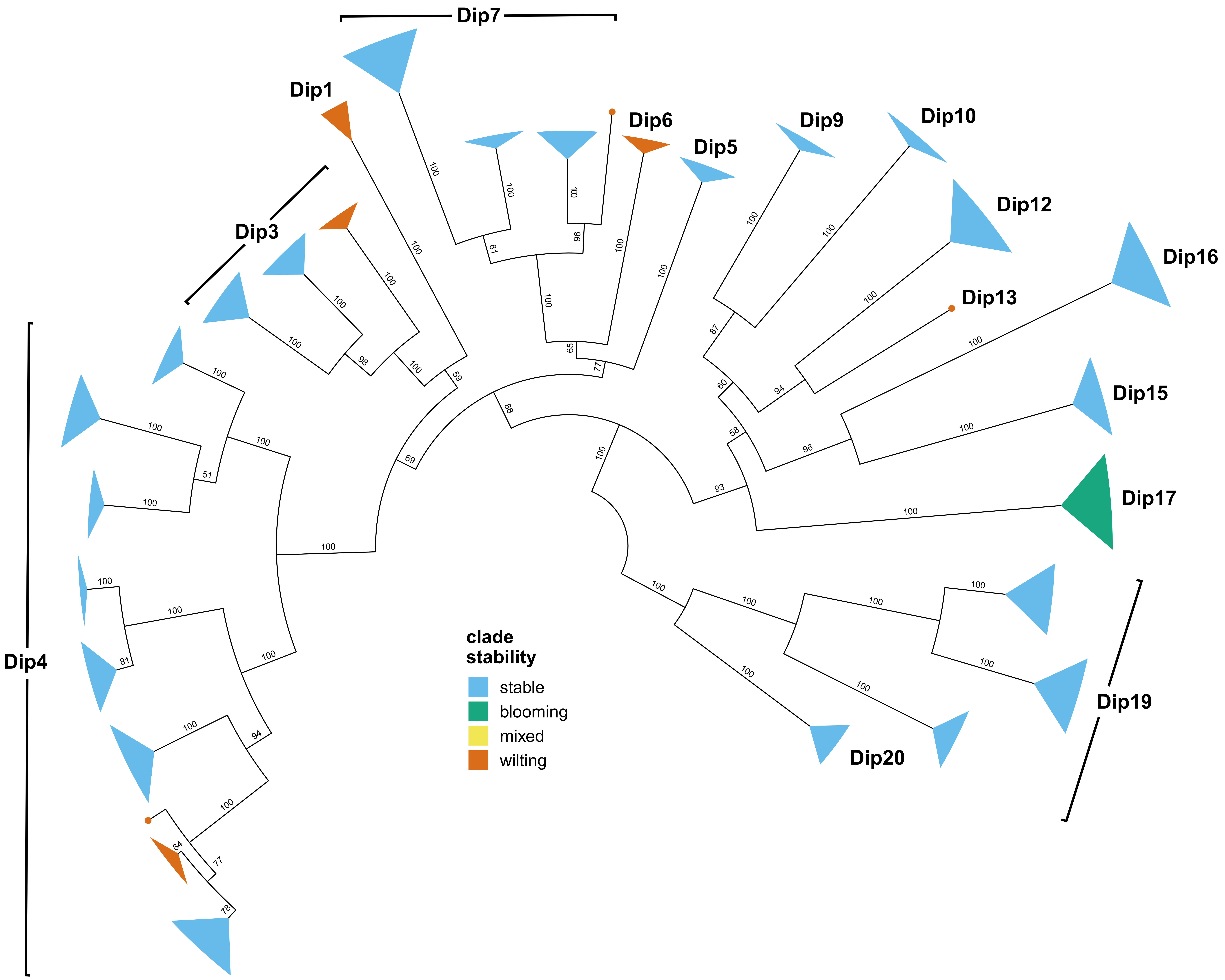

### Figure S11

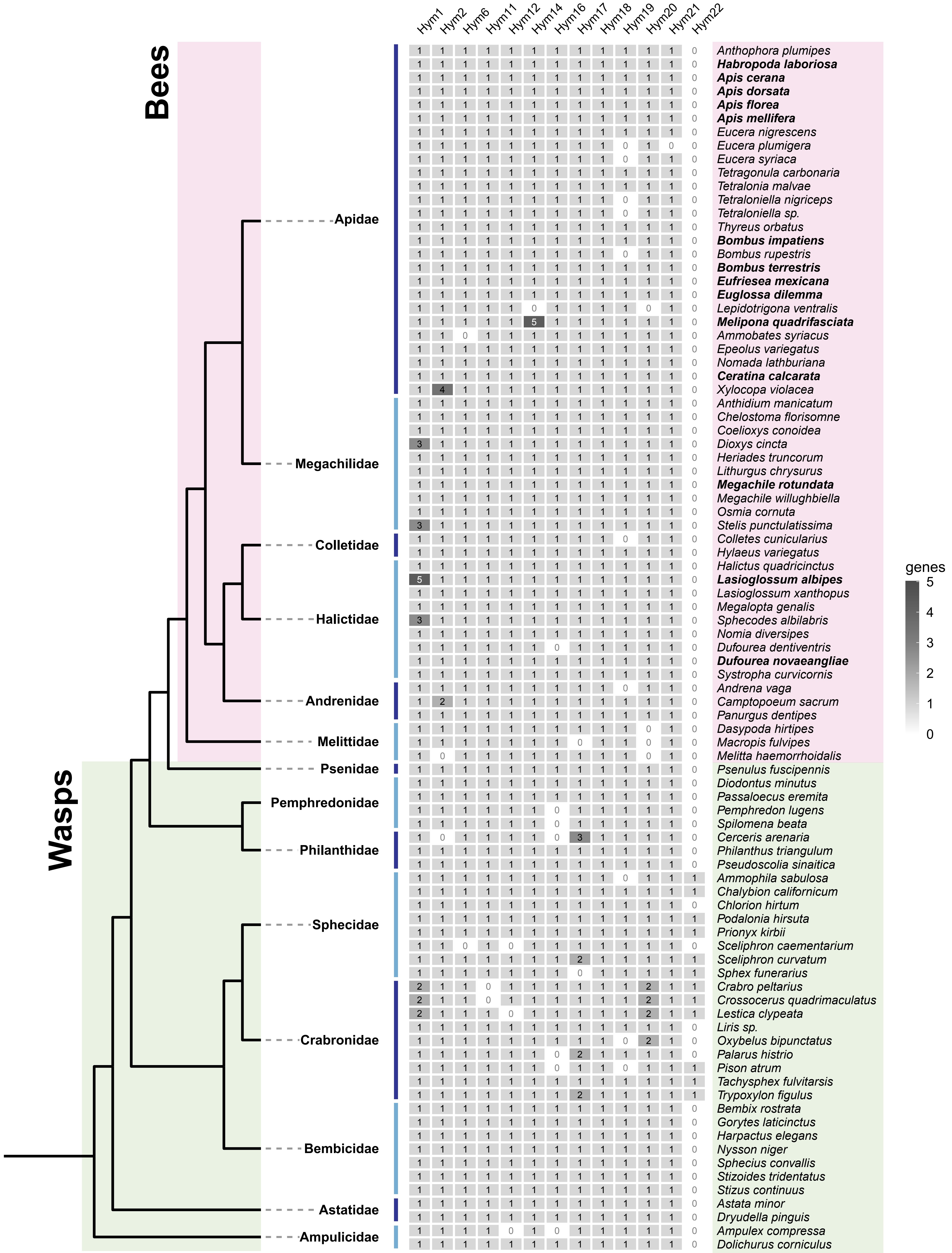
